# Xrp1 governs the stress response program to spliceosome dysfunction

**DOI:** 10.1101/2023.04.06.535851

**Authors:** Dimitrije Stanković, Luke Tain, Mirka Uhlirova

## Abstract

Co-transcriptional processing of nascent pre-mRNAs by the spliceosome is vital to regulating gene expression and maintaining genome integrity. Here, we show that the deficiency of functional U5 snRNPs in *Drosophila* imaginal cells causes extensive transcriptome remodeling and accumulation of highly mutagenic R-loops, triggering a robust stress response and cell cycle arrest. Despite compromised proliferative capacity, the U5 snRNP deficient cells increased protein translation and cell size, causing intra-organ growth disbalance before being gradually eliminated via apoptosis. We identify the Xrp1-Irbp18 heterodimer as the primary driver of transcriptional and cellular stress program downstream of U5 snRNP malfunction. Knockdown of *Xrp1* or *Irbp18* in U5 snRNP deficient cells attenuated JNK and p53 activity, restored normal cell cycle progression and growth, and inhibited cell death. Reducing Xrp1-Irbp18, however, did not rescue the splicing defects and the organismal lethality, highlighting the requirement of accurate splicing for cellular and tissue homeostasis. Our work provides novel insights into the crosstalk between splicing and the DNA damage response and defines the Xrp1-Irbp18 heterodimer as a critical sensor of spliceosome malfunction.

## INTRODUCTION

Pre-mRNA splicing catalyzed by the spliceosome is central to the regulation of gene expression in eukaryotes. Its physical and functional coupling with transcription ensures the production of cell- and condition-specific transcript variants required for development and the function of different cell types in multicellular organisms (Boumpas *et al*, 2022; Herzel *et al*, 2017; Tellier *et al*, 2020). Defects in the splicing machinery caused by mutations in spliceosome components or their dysregulation have been linked to tissue-, or even cell-type-specific pathologies in humans, and correlate with aging in a variety of organisms (Bhadra *et al*, 2020; Lee *et al*, 2016; Scotti & Swanson, 2016; Wood *et al*, 2021). The diversity of processes that depend on functional spliceosomes, and the phenotypes inflicted by spliceosome dysfunction, underscores the complexity of the molecular machine and highlights the need for in-depth understanding of its integral role in gene regulation in an *in vivo* context.

The spliceosome is a large ribonucleoprotein complex composed of five small nuclear ribonucleoprotein particles (snRNPs) each of which comprises specific uridine-rich small nuclear RNA (U snRNA) and a set of core and auxiliary proteins (Will & Luhrmann, 2011). In multicellular eukaryotes, most introns are removed by major (U2) spliceosomes, which contain U1, U2, U4/U6 and U5 snRNPs. The minor (U12) spliceosome, comprising U11, U12, Y4atac, U5 and, U6atac (Tarn & Steitz, 1996a; Tarn & Steitz, 1996b) processes less than 0.5% of introns (Moyer *et al*, 2020). The U5 snRNP is the only snRNP common to both spliceosome types, playing a vital role in spatial organization of the catalytic center, positioning of critical U snRNA and substrate exon residues, and timely activation of the spliceosome (Matera & Wang, 2014; Wahl *et al*, 2009; Will & Luhrmann, 2011). The U5 snRNP consists of U5 snRNA, Sm-protein ring, several accessory proteins, and core factors, including the pre-mRNA processing factor 8 (PRPF8/Prp8) (Grainger & Beggs, 2005). Despite its universal requirement for pre-mRNA splicing, mutations in Prp8 cause retina-specific disease, an autosomal dominant form of the Retinitis Pigmentosa type 13 (RP13) (Ruzickova & Stanek, 2017; Wood *et al*., 2021). The U5 snRNP joins the spliceosome as part of the U4/U6•U5 tri-snRNP, prior to which it undergoes a highly coordinated, multistep assembly process, that takes part in both the cytoplasm and the nucleus. The cytoplasmic phase requires an evolutionarily conserved protein Ecdysoneless (Ecd) (Claudius *et al*, 2014; Cloutier *et al*, 2017; Erkelenz *et al*, 2021; Malinova *et al*, 2017) that promotes U5 snRNP biogenesis through interactions with Prp8, the Sm ring protein SmD3, and U5 snRNA (Erkelenz *et al*., 2021). In the absence of Ecd, Prp8 is destabilized, and U5 snRNP biogenesis is stalled, resulting in apoptosis and eventually lethality in the *Drosophila* model (Claudius *et al*., 2014; Erkelenz *et al*., 2021). Similarly, as with malfunction of other core U5 snRNP components (Mordes *et al*, 2006; Rubio-Peña *et al*, 2015; Ruzickova & Stanek, 2017; Stankovic *et al*, 2020; Yang *et al*, 2021), Ecd deficiency causes genome-wide changes in gene expression and splicing, and has been linked to a diverse array of phenotypes, including reduced proliferation (Kim *et al*, 2009), aberrant nuclear mRNA export (Saleem *et al*, 2021), and activation of both unfolded protein- and oxidative stress-responses (Erkelenz *et al*., 2021; Olou *et al*, 2017). However, knowledge on the cascade of events and mechanisms that trigger or contribute to loss of cellular homeostasis, and ultimately death, remain incomplete.

Here, we exploit the genetic accessibility of the *Drosophila* model to elucidate the course of events arising from scarcity of the U5 snRNP inflicted by knockdown of the biogenesis factor Ecd or an RP13-associated mutation in Prp8. We identify Xrp1 and its dimerizing partner Irbp18 as the main drivers of transcriptional changes and cellular stress responses to U5 snRNP malfunction. While Xrp1/Irbp18 knockdown alleviated the harmful consequences of prolonged stress signaling it did not restore spliceosome fidelity, highlighting the primary role and requirement of U5 snRNP in maintenance of cellular homeostasis by controlling pre-mRNA splicing.

## RESULTS

### Spliceosome malfunction induces stress signaling and progressive apoptosis

Ecd has emerged as a crucial mediator of U5 snRNP biogenesis (Claudius *et al*., 2014; Cloutier *et al*., 2017; Erkelenz *et al*., 2021; Malinova *et al*., 2017). To characterize the sequence of events triggered by spliceosome dysfunction on the cellular and tissue level, we reduced Ecd function in the pouch region of the developing wing imaginal disc (WD) by expressing transgenic *ecd* RNAi (*UAS-ecd^RNAi^*), under the control of the *nubbin-Gal4* driver (Figure 1A) (hereafter referred to as *nub>*). Importantly, suppressing *ecd* expression in actively cycling imaginal cells of the WD pouch domain allowed normal development until the late 3^rd^ instar larval stage. However, in contrast to control larvae (*nub>/+*), the larval-to-pupal transition of *nub>ecd^RNAi^*animals was delayed, with the majority dying during metamorphosis. The few adults that managed to eclose lacked wing blades (Figure 1B-C).

**Figure 1.**
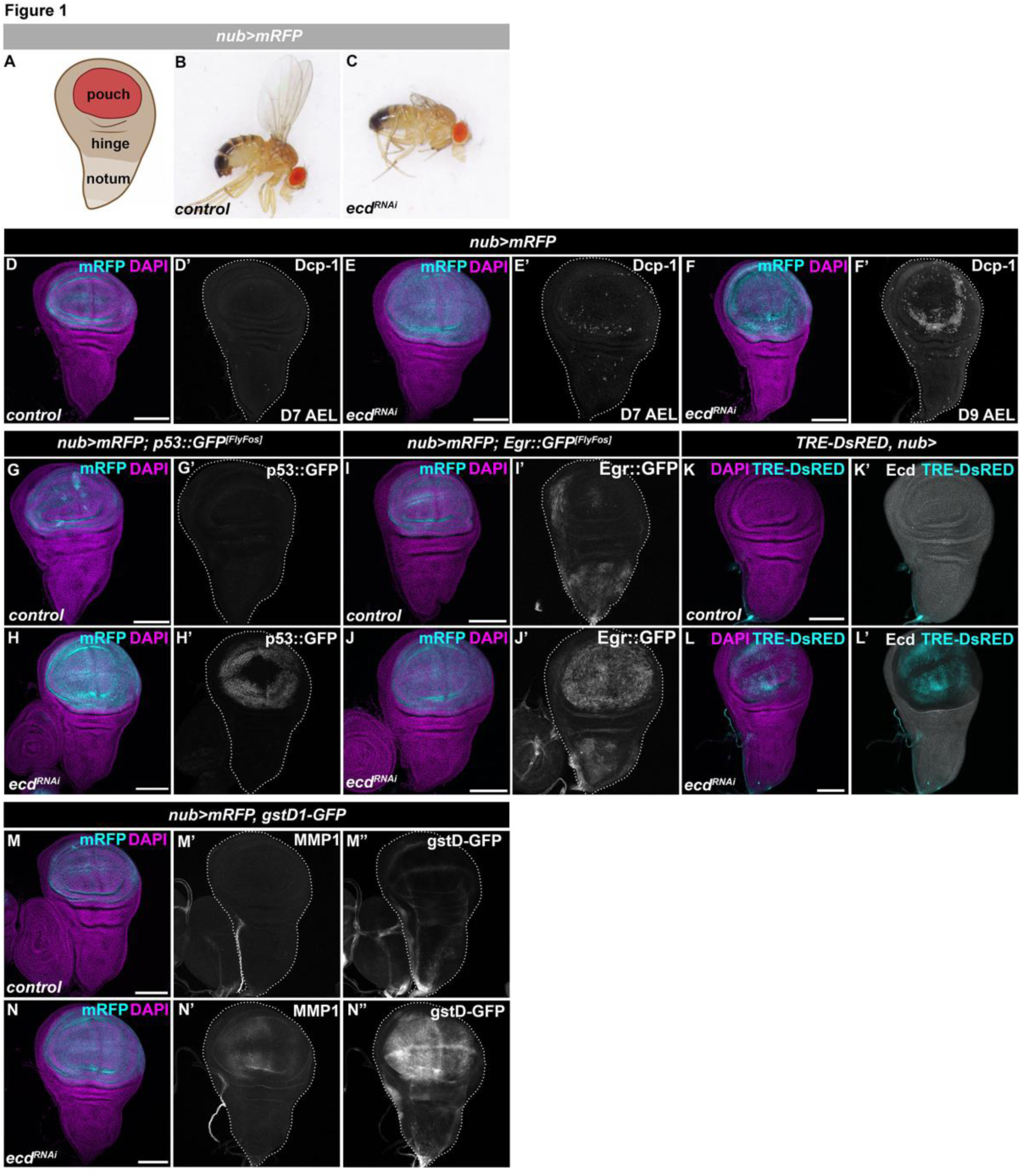
Ecd deficiency results in increased cell death and stress response signaling involving JNK and p53. **(A)** Schematic representation of the *Drosophila* wing imaginal disc comprising pouch, hinge, and notum. The *nub-Gal4* driver is active the pouch region highlighted by mRFP expression. **(B-C)** Representative image of control (*nub>mRFP*) (B) and a rare *nub>mRFP ecd^RNAi^*adult escaper male fly lacking the wing blade (C). **(D-F)** Immunostaining shows increased Dcp-1 signal in pouch region of *nub>mRFP ecd^RNAi^* WD on day 7 (E’) with further enhancement on day 9 AEL (F’), compared to D7 AEL control (*nub>mRFP*) WD (D’). Note, the Dcp-1 signal is not ubiquitous across the nubbin domain but enriched around the perimeter of the *nub>mRFP ecd^RNAi^* wing pouch (E’, F’). **(G-J)** Immunostaining shows increased levels of GFP-tagged p53 and Egr FlyFos transgenes in *nub>mRFP ecd^RNAi^* WDs at 7 days AEL (H, J) compared to control WDs (G, I). **(K-L)** JNK activity is increased while Ecd levels are reduced in pouch domain of *nub>ecd^RNAi^* (L) WD compared to control (*nub>*) WDs (K) as determined by immunostaining for *TRE-DsRED* transgenic reporter and Ecd protein. **(M-N)** Immunostaining shows increased levels of MMP1 (N’) and expression of a *GstD1-GFP* transgenic reporter (N’’) in pouch region of *nub>mRFP ecd^RNAi^* WDs compared to control WDs (M’, M’’). The confocal micrographs of WDs show maximum intensity projections of multiple confocal sections. DAPI and mRFP label nuclei and the nubbin domain, respectively. WDs are outlined by white dotted lines based on DAPI staining. Scale Bars: 100 μm. See also Supplementary Figure S1.

In agreement with previous studies (Claudius *et al*., 2014; Erkelenz *et al*., 2021; Redfern & Bownes, 1983) and the adult wingless phenotype (Figure 1C), Ecd deficiency caused progressive apoptosis within the pouch domain, as demonstrated by immunostaining of activated Dcp-1, the main effector caspase of *Drosophila* (Figure 1D-F and Supplementary Figure 1A-D). Compared to a moderate enrichment of apoptotic cells in *nub>ecd^RNAi^*WDs on day 7 AEL relative to control, large patches of Dcp-1 positive cells were observed in delayed 9-day old larvae (Figure 1D-F and Supplementary Figure 1A-D). In contrast to the limited cell death on day 7 AEL, wing pouch cells with reduced Ecd levels showed a strong activation of stress-induced signaling pathways and their downstream effectors. Using immunostaining of endogenous or a transgenic FlyFos-based tagged protein, we detected upregulation of p53 (Figure 1G-H and Supplementary Figure 1E-G), an evolutionarily conserved regulator of genome stability, cell survival, senescence, and apoptosis (Brodsky *et al*, 2004; Jin *et al*, 2000; Ollmann *et al*, 2000; Vousden & Prives, 2009). The enrichment of p53 protein in the *nub>ecd^RNAi^* wing pouch cells coincided with activation of the JNK signaling pathway, demonstrated by increased expression of a tumor necrosis factor (TNF) ligand Eiger (Egr) (Figure 1I, J), a primary upstream activator of JNK signaling (Igaki *et al*, 2002), the *TRE-DsRED* transgenic transcriptional reporter (Chatterjee & Bohmann, 2012) (Figure 1K, L), and the accumulation of a JNK-regulated target gene Matrix metalloprotease 1 (MMP1) (Uhlirova & Bohmann, 2006) (Figure 1M’, N’). Moreover, Ecd deficient cells upregulated *gstD1-GFP* (Figure 1M”, N”), a transcriptional reporter sensitive to changes in cellular redox homeostasis and proteotoxic stress (Brown *et al*, 2021; Langton *et al*, 2021; Sykiotis & Bohmann, 2008). These results highlight activation of p53- and JNK-mediated signaling among the early responses to reduced Ecd levels. Furthermore, *gstD1-GFP* and p53 upregulation was also observed in imaginal disc cells overexpressing the pathogenic RP13-linked mutant variant of Prp8 protein (*Drosophila* Prp8^[S2178F]^/human PRPF8^[S2118F]^) (Supplementary Figure 1H-M), suggesting a shared response to U5 snRNP malfunction, inflicted either by scarcity of the biogenesis factor or mutation in the core splicing factor. Importantly, blocking apoptosis by overexpression of p35, a pan-caspase inhibitor, prevented physical cell elimination (Supplementary Figure 1A-D) but did not restore the cellular homeostasis (Supplementary Figure 1E-G, J-M) required for normal adult appendage development. Together, this suggests that rather than acting as robust apoptotic inducers that ensure rapid cell elimination, p53 and JNK may work in concert to mount cytoprotective responses promoting cellular senescence-like state to delay cell death.

### Ecd scarcity induces signs of DNA damage and increases RNase H1 binding to DNA

One of the emerging hallmarks of spliceosome dysfunction is DNA damage and loss of genome integrity (Li & Manley, 2005; Naro *et al*, 2015; Paulsen *et al*, 2009; Tresini *et al*, 2015). This damage manifests as DNA double-strand breaks (DSBs) and the accumulation of RNA:DNA hybrid structures, known as R loops, in which the nascent transcript anneals to the DNA template strand (Aguilera & García-Muse, 2012). The fact that loss of Ecd interferes with U5 snRNP biogenesis and induces the major DNA damage sensor p53 indicated that Ecd deficient cells could suffer from DNA damage and ectopic R-loop formation. Indeed, immunostaining revealed increased number of pH2Av foci, marking DSBs, in Ecd depleted cells of the wing pouch compared to the rest of the tissue and control WDs (Figure 2A-C). Furthermore, Ecd deficient cells showed upregulation of the Inverted repeat-binding protein (Irbp) (Figure 2D, E), the *Drosophila* homolog of the human XRCC6 (Ku70 protein), which forms heterodimers with Ku80 to participate in DSB repair via nonhomologous end joining (NHEJ) (Beall *et al*, 1994; Jacoby & Wensink, 1994). Importantly, strong enrichment of Irbp::GFP was also observed in wing pouch cells overexpressing the pathogenic Prp8^[S2178F]^ protein (Supplementary Figure 1L, M). In addition, we detected increased expression of *RNase H1* (*rnh1*) mRNA, an enzyme that is recruited to, and degrades the RNA moiety of R-loops (Crossley *et al*, 2019), in WDs bearing Ecd deficient cells (Figure 2F). To map genome-wide distribution of RNase H1, as a proxy for R-loop formation, we performed Targeted DamID (TaDa) (Southall *et al*, 2013) on *control* and *ecd^1^* homozygous mutant WDs expressing Dam-RNase H1 fusion protein in the nubbin domain (Figure 2G). The Dam only expressing control and *ecd^1^* mutant WD samples served as normalization controls for the experiment. In both control and *ecd^1^* mutant imaginal tissue, the majority of Dam-RNase H1 peaks were associated within promoter-proximal regions (Figure 2H). However, *ecd^1^* mutants showed an increase in both the mean intensity and total number of peaks (Figure 2I, J). Moreover, the analysis further revealed a 2.45 times greater number of genes associated with increased Dam-RNase H1 binding in *ecd^1^*mutants than in controls (Figure 2K and Supplementary Dataset S1). In summary, these data show that spliceosome malfunction inflicted by Ecd loss results in DNA damage. Increased expression and binding of RNase H1 in Ecd deficient cells further suggest the aberrant formation of R-loops that may contribute to the dysregulation of gene expression and genome instability (Aguilera & García-Muse, 2012; Niehrs & Luke, 2020).

**Figure 2.**
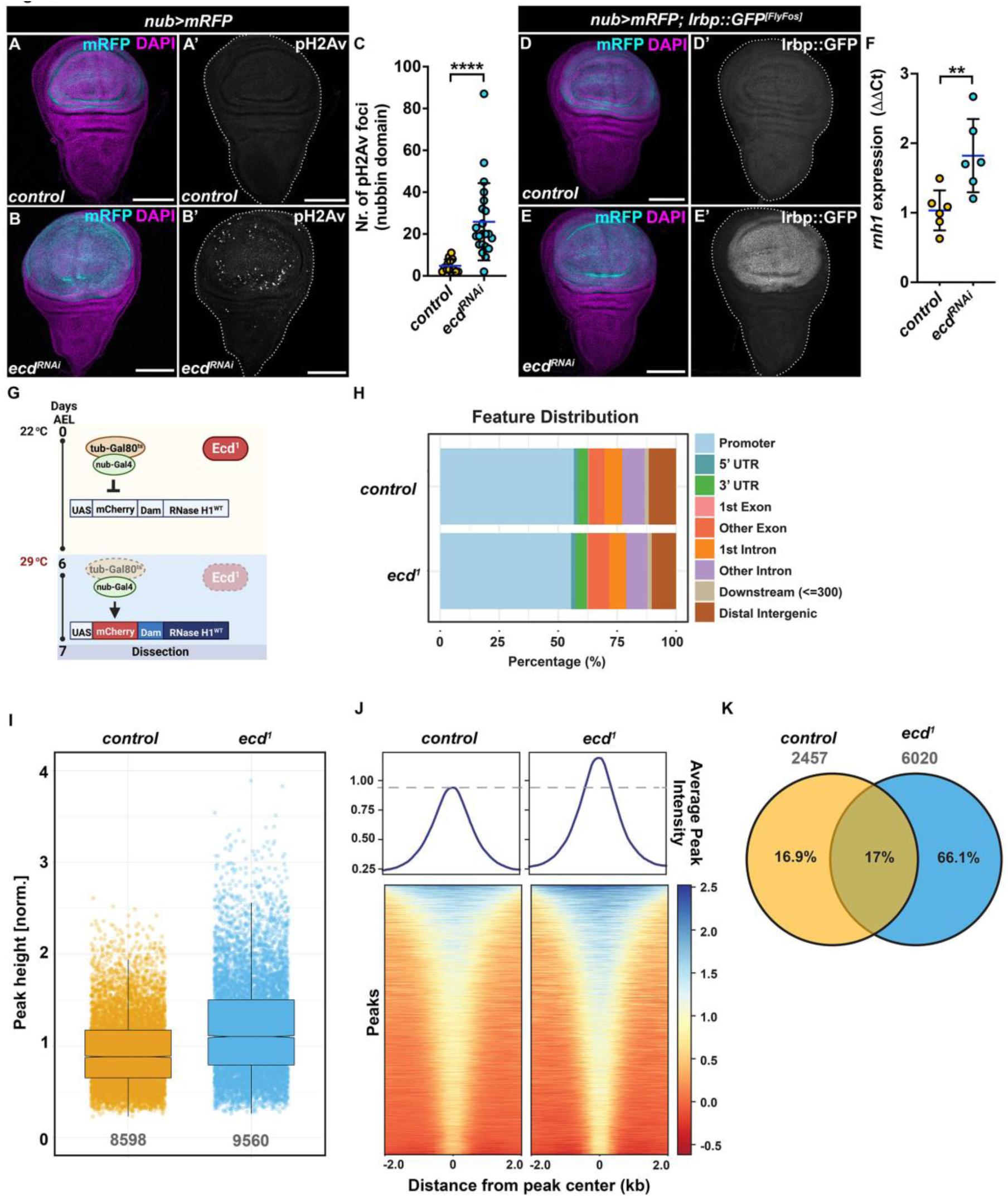
DNA damage induced by Ecd loss correlates with increased RNase H1 binding to DNA. **(A-C)** Representative confocal micrographs (A, B) and quantification (C) show increased in phosphorylation of H2Av in wing pouch of *nub>mRFP ecd^RNAi^* (B) (n = 22) compared to *control* WDs (A) (n = 13) at day 7 AEL as visualized with an antibody against phosphorylated histone variant H2Av (pH2Av). Scale Bars: 100 μm. For quantification (C), the number of pH2Av foci were quantified within the nubbin domain. Unpaired nonparametric two-tailed Mann-Whitney test was used to calculate p-values. **(D-E)** Wing pouch-specific knockdown of *ecd* (*nub>mRFP ecd^RNAi^*) increases expression of Irbp::GFP FlyFos reporter (E’) compared to control WDs (D’) at day 7 AEL. Scale Bars: 100 μm. **(F)** Fold change in expression of endogenous *rnh1* transcript in *nub>mRFP ecd^RNAi^* WDs relative to control as determined by RT-qPCR analysis (ΔΔCt), n = 6. Levels of TATA binding protein (*tbp*) transcripts were used for normalization. Unpaired Student’s t-test assuming equal variances was used to determine statistical significance. **(G)** Schematic representation of the TaDa experimental design taking advantage of the temperature-sensitive *ecd^1^*mutant allele and an α-tubulin promoter-driven transcriptional repressor Gal80^ts^ (tub-Gal80^ts^). Gal80^ts^ blocks expression of *UAS-mCherry-Dam-RNase H1^WT^* or *UAS-mCherry-Dam* fusion construct by the *nub-Gal4* driver in *ecd^1^* homozygous mutant or control larvae when kept at the permissive temperature (25 °C). On day 6 AEL, the larvae were transferred to a non-permissive temperature (29 °C) to simultaneously abolish the function of Ecd and induce expression of the UAS-Dam transgenes. WDs samples were dissected on day 7 AEL. **(H)** Distribution of significant Dam-RNase H1 peaks in *control* and *ecd^1^* conditions, in relation to different genomic features expressed as percentage of all features. The feature distribution was obtained using the *ChipSeeker* R package (Yu *et al*, 2015). **(I)** Scatter plot of intensities of all significant Dam-RNase H1 peaks in *control* and *ecd^1^* WDs. The summary of the data is shown as a boxplot and interquartile range (IQR). Whiskers show 1.5*IQR. **(J)** Dam-RNase H1 peak intensity profiles across a 4 kb region centered on all significant peaks detected in both control and *ecd^1^* WDs. The heatmap color and profile plots show the higher average Dam-RNase H1 peak intensity across all significant peaks in *ecd^1^* WDs. **(K)** Venn diagram shows the overlap between genes in proximity of Dam-RNase H1 binding events that are higher in either control or *ecd^1^* WDs. The majority of GATC fragments throughout the genome are characterized by higher Dam-RNase H1 peak intensities in *ecd^1^* compared to the control. The overlap between two groups represents gene-proximal local redistribution of Dam-RNase H1 binding events. The confocal micrographs of WDs show maximum intensity projections of multiple confocal sections unless stated otherwise. DAPI and mRFP label nuclei and the nubbin domain, respectively. WDs are outlined by white dotted lines based on DAPI staining. Data in charts represent means ± SD, \*\**p* < 0.01, \*\*\*\**p* < 0.0001. See also Supplementary Figure S1 and Supplementary Dataset S1.

### *Ecd* deficiency results in G2-phase cell cycle arrest and aberrant growth

In the physiological context, accumulation of DNA damage triggers DNA-damage checkpoints that cause transient or permanent cell cycle arrest depending on the extent of damage, and the capacity of cell to repair it. To determine the replicative and proliferative capacity of WD cells with compromised Ecd function, we performed EdU labelling along with immunostaining for the mitotic marker, phosphorylated Histone H3 (pH3). We observed significant reduction in the number of EdU- and pH3-positive cells in the nubbin compartment of the *nub>ecd^RNAi^*WDs compared to controls (Figure 3A-E), indicative of cell cycle stalling or arrest. However, the decrease in mitosis also occurred in hinge and notum region of *nub>ecd^RNAi^* WDs (Figure 3E), suggesting possible intra-organ and systemic regulation of proliferation (Boulan *et al*, 2019; Sanchez *et al*, 2019). To define the precise cell cycle stage in which the *nub>ecd^RNAi^* WD cells arrest, we utilized the transgenic fly ubi-FUCCI system (Fluorescent Ubiquitination-based Cell Cycle Indicator), based on fluorochrome-tagged degrons from the Cyclin B (mRFP1-CycB1-266) and E2F1 (GFP-E2F11-230) proteins, expressed from the ubiquitin promoter (Zielke *et al*, 2014). Under normal conditions CycB is targeted for destruction by the Anaphase-promoting Complex (APC/C) during mitosis and early G1 phase, while E2F1 is degraded by CRL4^Cdt2-^ during S phase. Accumulation of both mRFP1-CycB_1-266_ and GFP-E2F1_1-230_ marks cells in G2, a phase associated with cell growth and elevated protein synthesis (Figure 3F). Given the delayed apoptosis and high JNK activity that has been shown to promote G2-phase stalling (Cosolo *et al*, 2019), we hypothesized that *ecd* deficient cells would be positive for both fluorescent degrons. Indeed, in contrast to control WDs, which comprised cells in all phases of the cell cycle in a developmentally controlled cell-cycle pattern (Figure 3G), the majority of *nub>ecd^RNAi^* WD cells were arrested in G2 phase (Figure 3H). Despite reduced mitotic activity (Figure 3C-E), Ecd deficient cells retained their growth potential and grew significantly larger compared control *nub>* WD cells (Figure 3I-K). This manifested as an increased size of the nubbin domain of *nub>ecd^RNAi^* WDs relative to the combined size of the hinge and notum (Figure 3L). Interestingly, *ecd* knockdown cells also contained larger nucleoli (Figure 3M-O), as determined by immunostaining of Fibrillarin, a component of C/D box small nucleolar ribonucleoprotein (snoRNP) particles (Rodriguez-Corona *et al*, 2015). As changes in nucleoli and cell size have been correlated with the level of ribosomal biogenesis/translation (Buchwalter & Hetzer, 2017; Wu *et al*, 2022), we asked if *ecd* silencing altered the rate of protein synthesis using puromycin-based labelling (Aviner, 2020). Intriguingly, immunostaining of WDs following a pulse treatment of puromycin showed significantly higher signal in Ecd-depleted wing pouch cells compared to the rest of the tissue and to control WDs (Figure 3P-R). Together, these results demonstrate that the activation of stress signaling in Ecd deficient cells correlates with their arrest in the G2 phase of the cell cycle and attenuation of mitotic activity within the wing pouch, but also in neighboring parts of the tissue. Despite genomic instability, the G2 arrested Ecd deficient cells increase in size, likely due to elevated translational activity, which contributes to intra-organ growth disbalance affecting organ proportions.

**Figure 3.**
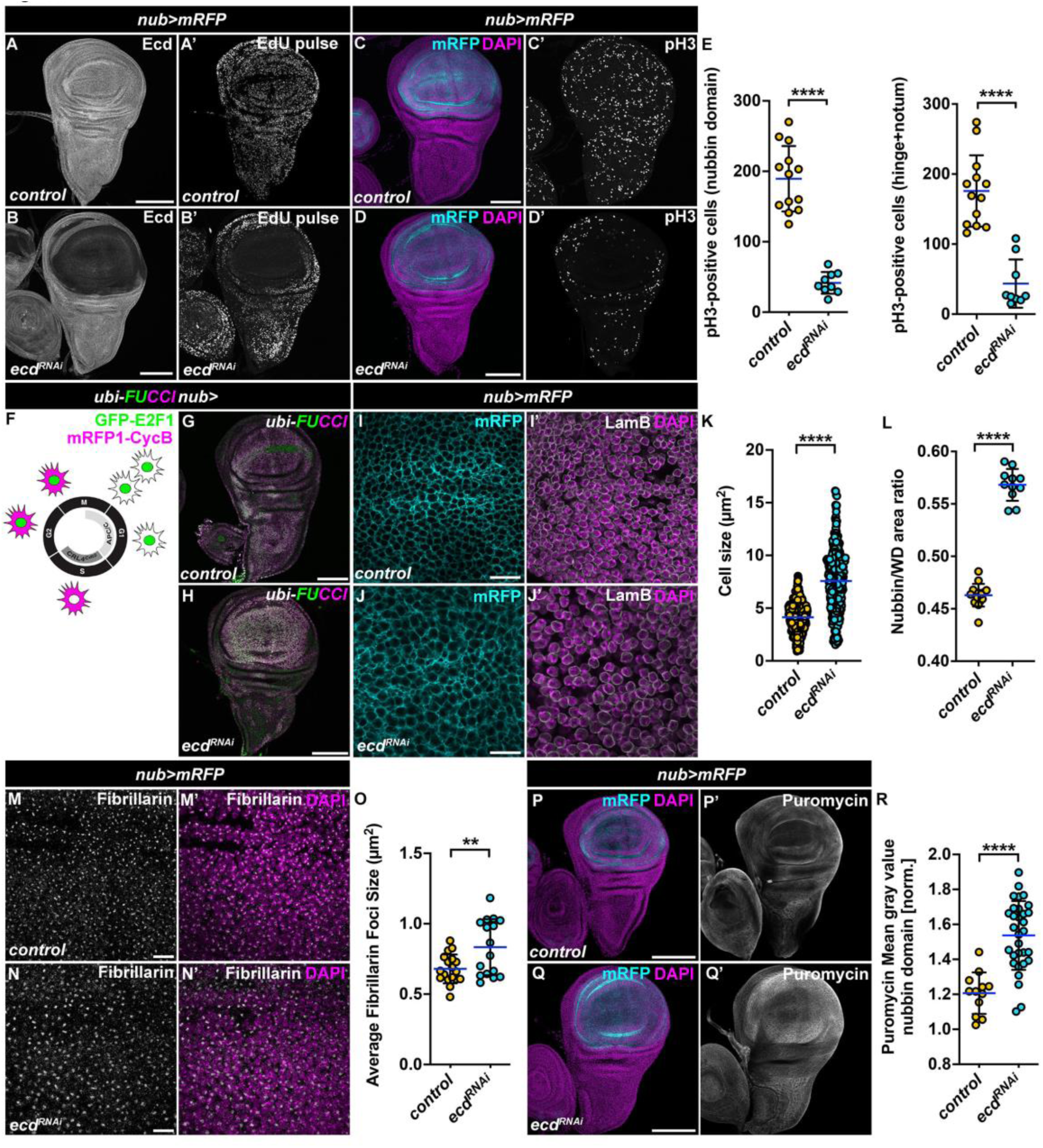
Cell cycle arrest caused by reduced Ecd is accompanied by increased cell growth and translation. **(A-B)** *In vivo* EdU incorporation is reduced in *nub>mRFP ecd^RNAi^* WDs (B’) compared to control (A’). Immunostaining with an Ecd-specific antibody shows efficient reduction of Ecd protein in the pouch region of WDs following overexpression of an *UAS-ecd^RNAi^*transgene relative to the rest of the tissue (B) and control WD (A). Scale Bars: 100 μm. **(C-E)** Immunostaining of WDs 7-days AEL with an anti-pH3 antibody revealed overall reduced mitotic activity in *nub>mRFP ecd^RNAi^* WD (D) compared to controls (C). Scale Bars: 100 μm. Quantification of pH3-positive cells in the nubbin domain and hinge + notum regions in control (n = 13) and *nub>mRFP ecd^RNAi^* (n = 9) WDs confirms negative impact, cell autonomous and non-autonomous, of *ecd* knockdown on cell proliferation (E). An unpaired Student’s t-test assuming equal variances was used to determine statistical significance. **(F-H)** Schematic of the fly ubi-FUCCI (ubi-GFP-E2F11-230 and ubi-mRFP1-NLS-CycB1-266) cell cycle reporter shows G1 (GFP), S (RFP) and G2/M (GFP and RFP) phases of the cell cycle (F). Compared to control (*ubi-FUCCI, nub>*) (G), many Ecd deficient cells in the pouch region of *ubi-FUCCI, nub>ecd^RNAi^*WD arrest in G2 phase of the cell cycle (GFP and RFP, yellow/white overlay) (H). **(I-K)** High magnification images of the nubbin domain of control (I) and *nub>mRFP ecd^RNAi^* WDs (J) at day 7 AEL. DAPI and Lamin B mark the nucleus and nuclear lamina, respectively. The images are projections of 2-3 consecutive confocal sections. Scale Bars: 10 μm. mRFP labelling of cell membranes was used for quantification of cell sizes (K), see Materials and Methods, (n = 7 WDs per genotype). **(L)** Wing pouch-specific knockdown of *ecd* (*nub>mRFP ecd^RNAi^*) (n = 14) leads to wing pouch overgrowth in relation to the hinge and notum of the WDs compared to control (*nub>mRFP*) (n = 13). The data represent a ratio of the nubbin domain and the entire WD at 7 days AEL. An unpaired two-tailed Student’s t-test assuming equal variances was used to determine statistical significance. **(M-O)** Representative images (M, N) and quantification (O) show enlarged nucleoli in wing pouch cells of *nub>mRFP ecd^RNAi^* WDs (n = 16) (N) relative to control (M) (n = 18) at day 7 AEL. Nucleoli were visualized by immunostaining with an anti-Fibrillarin antibody. DAPI labels nuclei. Scale Bars: 20 μm. Quantification shows the average nucleolar size per replicate in both conditions. A two-tailed Student’s t-test assuming equal variances was used to determine statistical significance. **(P-R)** Representative confocal micrographs (P, Q) and quantification (R) show increased puromycin incorporation (20-minutes pulse) in wing pouch of *nub>mRFP ecd^RNAi^* (n = 31) (Q) compared to control WDs (P) (n = 12) at day 7 AEL as visualized with an anti-puromycin antibody. Scale Bars: 100 μm. An unpaired Student’s t-test assuming equal variances was used to determine statistical significance. The images are projections of multiple confocal sections of WDs at day 7 AEL unless stated otherwise. DAPI and mRFP label nuclei and the nubbin domain, respectively. Data in charts represent means ± SD, \*\**p* < 0.01, \*\*\*\**p* < 0.0001. See also Supplementary Figure S3.

### Transcriptome profiling of *ecd* deficient cells highlights upregulation of translation, stress and DNA damage responses

To define the molecular network orchestrating the responses to Ecd scarcity, we profiled the transcriptome of control (*nub>*) and *nub>ecd^RNAi^* WDs using mRNA-seq (Supplementary Figure S2A-B). In total, we detected transcripts associated to 17,621 genes (Supplementary Dataset S2). Most differentially expressed genes (DEGs) in *nub>ecd^RNAi^* were protein coding genes (Figure 4A, B) of which 2912 were downregulated and 3166 upregulated compared to controls (uniquely identified or overlapping genes, *pAdj* < 0.05) (Figure 4A and Supplementary Dataset S2). The second most numerous and regulated transcript biotype corresponded to transposable elements (TEs) (Figure 4B). In total, we detected transcripts from 399 TEs, 273 of which were significantly up- and 126 down-regulated in *ecd^RNAi^* compared to controls (Supplementary Figure S2C and Supplementary Dataset S2). The gene ontology (GO) enrichment analysis of DEGs significantly upregulated in *nub>ecd^RNAi^* WDs showed a clear enrichment for functional categories associated with translation, ribosome and peptide biosynthesis, DNA damage and repair, autophagy, and glutathione metabolism (Figure 4C). In contrast, significantly downregulated genes were overrepresented for a diverse array of biological processes and pathways, including proteasome and spliceosome, and signaling pathways vital to imaginal disc and epithelium development, tissue morphogenesis and neurogenesis (Supplementary Figure S2D).

**Figure 4.**
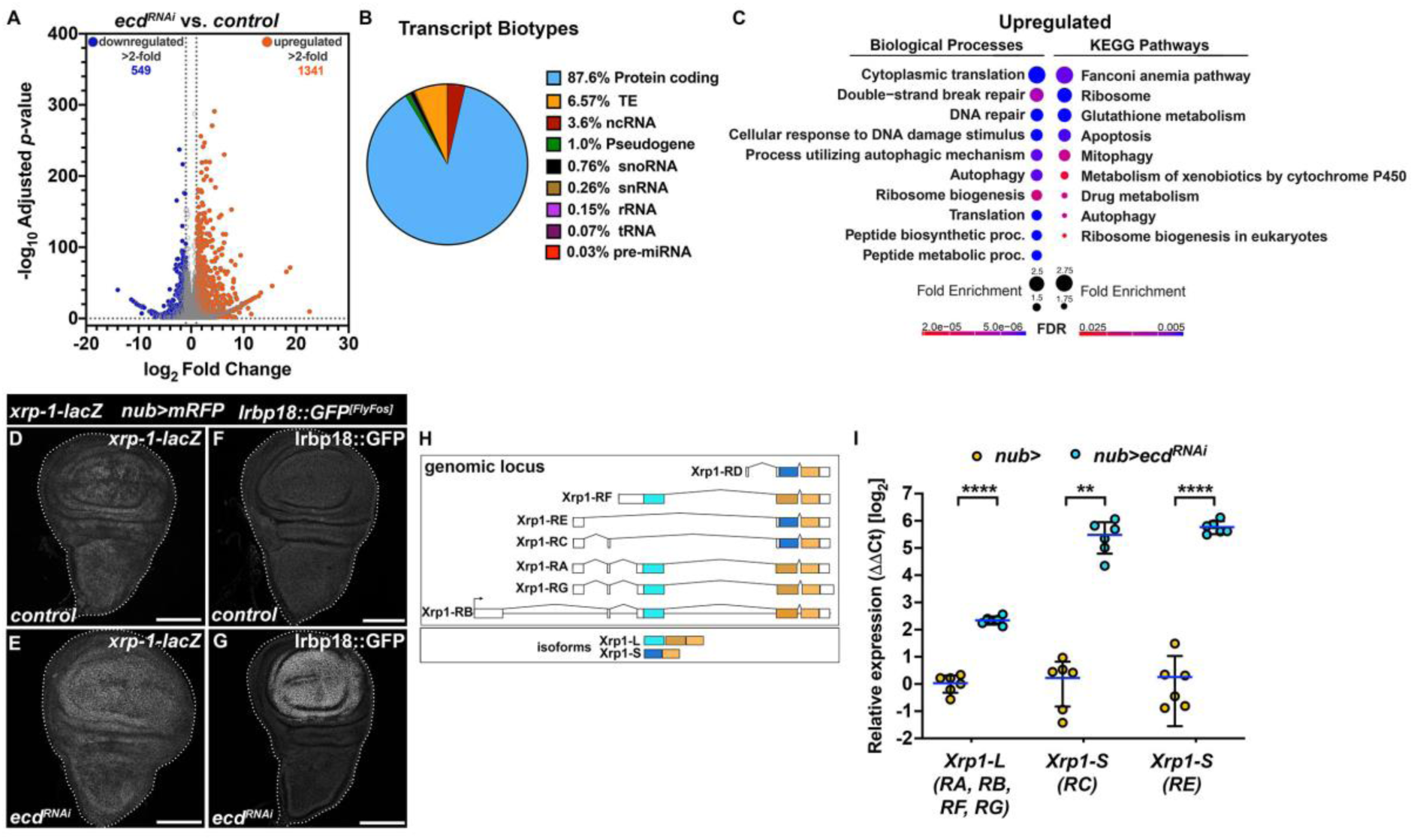
The transcriptome is remodeled in response to reduced Ecd levels, including upregulation of Xrp1 and Irbp18 C/EBP class of bZIP transcription factors. **(A)** The volcano plot depicts the significantly regulated genes in *nub>mRFP ecd^RNAi^* WDs compared to controls (*nub>mRFP*) detected by RNA sequencing. Genes up- and down-regulated >2-fold are shown in orange and blue, respectively. **(B)** Pie chart shows the proportion of each transcript biotype detected as significantly regulated (*pAdj* < 0.05) in response to reduced Ecd levels in *nub>mRFP ecd^RNAi^* WDs compared to controls (*nub>mRFP*). **(C)** GO enrichment plot for significantly upregulated genes (*pAdj* < 0.05) in *nub>mRFP, ecd^RNAi^* WDs compared to controls (*nub>mRFP*). Plots show fold enrichment of the top 10 significant functional GO terms. **(D-G)** Immunostaining shows elevated expression of *xrp1-lacZ* and Irbp18::GFP FlyFos reporter constructs *nub>mRFP ecd^RNAi^* (E, G) compared to control WDs (D, F) at day 7 AEL. The images are projections of multiple confocal sections. WDs are outlined by white dotted lines based on DAPI staining. Scale Bars: 100 μm. **(H-I)** Schematic of the *xrp1* genomic locus encoding 7 different transcripts giving rise to long (Xrp1-L) and short (Xrp1-S) protein isoforms (H). RT-qPCR analysis revealed strong upregulation of *xrp1* transcripts in *nub>mRFP ecd^RNAi^* WDs relative to control samples (*nub>mRFP*), particularly RC and RE coding for Xrp1-S protein isoform (I). Levels of *tbp* transcript were used for normalization. Unpaired Student’s t-test assuming equal variances was used to determine statistical significance. Data represent means ± SD, n = 6, \*\**p* < 0.01, \*\*\*\**p* < 0.0001. Statistical significance of differences between isoform expression was determined using two-way ANOVA with Sidak’s correction for multiple comparisons. See also Supplementary Figure S2 and Supplementary Dataset S2.

To identify the potential drivers of the transcriptional changes in Ecd deficient cells, we analyzed the expression of 757 known or predicted transcription factors (Adryan & Teichmann, 2006) (https://www.mrc-lmb.cam.ac.uk/genomes/FlyTF/). Given the substantial changes to the transcriptome, unsurprisingly, over half of the examined TFs (385) were differentially expressed in response to Ecd silencing using *pAdj* < 0.05 as a cut off (Supplementary Dataset S2). Of the 181 TFs whose expression increased, Structure specific recognition protein (Ssrp), Inverted repeat binding protein 18 (Irbp18), and Xrp1, were the three most significantly regulated. Intriguingly, all three proteins have been associated with DNA repair and/or maintenance of genome stability (Akdemir *et al*, 2007; Francis *et al*, 2016; Hsu *et al*, 1993). While Ssrp is a high mobility group (HMG) box transcription factor controlling chromatin organization as part of the Facilitates Chromatin Transcription (FACT) complex (Belotserkovskaya *et al*, 2003; Shimojima *et al*, 2003). Xrp1 and Irbp18 are basic region leucine zipper (bZIP) proteins of the CCAAT/enhancer-binding protein (C/EBP) family whose heterodimers have been shown critical for repairing DSBs following P-element transposition (Akdemir *et al*., 2007; Francis *et al*., 2016). Recently, Xrp1-Irbp18 have been described as key regulators of proteotoxic stress responses controlling PERK-dependent expression of antioxidant genes and elimination of looser cells during cell competition (Blanco *et al*, 2020; Brown *et al*., 2021; Langton *et al*., 2021).

To confirm the increased expression of Xrp1 and Irbp18 in Ecd deficient cells detected in our transcriptomic analysis, we utilized a *xrp1-lacZ* transcriptional reporter and an *Irbp18::GFP* FlyFos transgenic line, respectively. Immunostaining showed a clear induction of Xrp1 and Irbp18 expression in wing pouch cells of the *nub>ecd^RNAi^*WDs relative to the rest of the tissue and control WDs (Figure 4D-G). Moreover, the RT-qPCR analysis revealed that of the multiple functional *Xrp1* isoforms (Figure 4H) transcripts encoding RC and RE short isoforms were upregulated to a greater extent than longer isoforms in response to *ecd* knockdown relative to control (Figure 4I).

In summary, our genome-wide RNA profiling revealed a high degree of transcriptional remodeling in wing imaginal tissue in response to *ecd* loss. The molecular signature determined by our bioinformatic analysis is in marked agreement with the experimental findings described above, both highlighting a targeted upregulation of the translational machinery, stress response, and DNA damage/repair responses among the key processes affected by Ecd deficiency. The upregulation of Xrp1 and Irbp18 further suggested that the two transcription factors could act as the major mediators of the responses and phenotypes in Ecd-depleted cells.

### Xrp1-Irbp18 drives the stress and growth responses in spliceosome compromised cells

To determine the requirement of Xrp1 and Irbp18 to Ecd deficient phenotypes, we silenced either of the two TFs in wing imaginal cells lacking Ecd. Simultaneous knockdown of Xrp1 abolished the increase in p53, Egr, MMP1, and GstD1 expression (Figure 5A-I), but also apoptosis and DNA damage, observed in *nub>ecd^RNAi^* WDs as demonstrated by immunostaining for Dcp1 and pH2Av, respectively (Figure 5J-L). It also restored the number of pH3 positive cells to control levels (Figure 5M-O), suppressed the G2 cell cycle arrest (Figure 5P, Q), reduced the protein synthesis rate (Figure 5R-T), reverted cell size to control levels (Figure 5U), and prevented wing pouch overgrowth (Figure 5V). Importantly, inhibiting Irbp18, which upregulation in Ecd deficient cells largely depends on Xrp1 (Supplementary Figure S3A-D), recapitulated the effect of Xrp1 loss (Supplementary Figure S3E-P). These results demonstrate a pivotal role for Xrp1 in activating the canonical JNK and p53 signaling and regulating downstream cellular responses to Ecd scarcity, including cell proliferation, cellular translation and growth, DNA damage, and apoptosis. Furthermore, our analysis suggests Xrp1 likely acts in concert with its known heterodimerizing partner Irbp18 to drive the features of cellular senescence.

**Figure 5.**
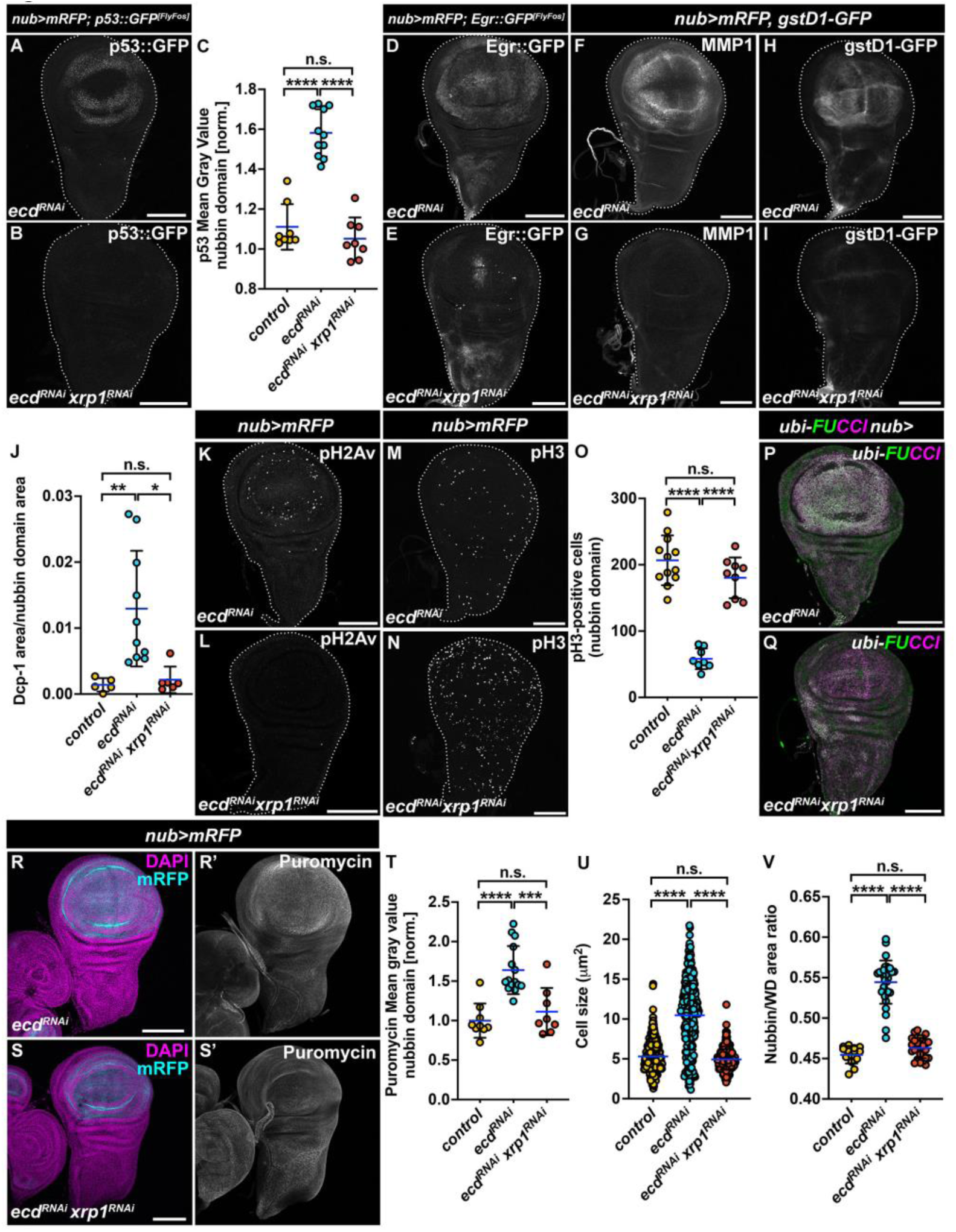
Xrp1 controls stress responses in Ecd deficient cells but is not responsible for lethality. **(A-J)** Immunostaining shows that the increased p53::GFP (A), Egr::GFP (D), MMP1 (F), and gstD1-GFP (H) expression in *nub>mRFP ecd^RNAi^* WDs is abolished by simultaneous knockdown of *xrp1* (B, E, G, I), while Dcp-1 levels are reverted to those in control WDs. (J) Quantification of p53::GFP signal in the nubbin domain of control (*nub>mRFP*), *nub>ecd^RNAi^* and *nub>ecd^RNAi^ xrp1^RNAi^*WDs (C). One-way ANOVA with Tukey’s multiple comparisons test was used to calculate *p*-values. **(K-L)** Phosphorylation of H2Av is Xrp1-dependent, as knockdown of *xrp1* leads to the loss of fluorescent signal corresponding to phosphorylated H2Av (pH2Av) in Ecd deficient (*nub>mRFP ecd^RNAi^ xrp1^RNAi^*) WDs. **(N-O)** Representative confocal micrographs (M, N) and quantification (O) show reduced pH3 signal in *nub>ecd^RNAi^* WDs (n = 9) that is normalized to control counts (n = 12) when *xrp1^RNAi^* is co-expressed with *ecd^RNAi^*(n = 9). One-way ANOVA with Tukey’s multiple comparisons test was used to calculate *p*-values for quantification of pH3-positive cells in the nubbin domain. **(P-Q)** G2 phase cell cycle arrest of wing pouch cells induced by Ecd deficiency (P) is reverted in *nub>ecd^RNAi^ xrp1^RNAi^*WDs (Q) as determined by immunostaining of ubi-FUCCI cell cycle reporter. **(R-T)** Representative confocal micrographs of WDs 7-days AEL stained with the anti-puromycin antibody (R, S) and fluorescent signal quantification (T) show that *xrp1* silencing (*nub>mRFP ecd^RNAi^ xrp1^RNAi^*) (n= 8) reverts the increased translation rates caused by *ecd* knockdown (*nub>mRFP ecd^RNAi^*(n = 15) WDs (T) to control levels (n = 9). One-way ANOVA with Tukey’s multiple comparisons test was used to calculate *p*-values. **(U, V)** Wing pouch-specific knockdown of *xrp1* using *nub>mRFP* driver completely abolishes increase in cell size (U) and overgrowth of the nubbin domain (V) caused by Ecd scarcity. Membrane-targeted mRFP signal was used to measure cell sizes localized on diagonal in the nubbin domain of control (*nub>mRFP*, n = 7), *nub>mRFP ecd^RNAi^* (n = 7), *nub>mRFP ecd^RNAi^ xrp1^RNAi^* (n= 7) WDs (U). A ratio of the nubbin domain and the entire WD were determined based on the mRFP and DAPI signals (V). Control (*nub>mRFP*, n = 14), *nub>mRFP ecd^RNAi^* (n = 31), *nub>mRFP ecd^RNAi^ xrp1^RNAi^*(n= 26). One-way ANOVA with Tukey’s multiple comparisons test was used to calculate *p*-values. The images are projections of multiple confocal sections of WDs 7-days AEL. DAPI and mRFP label nuclei and the nubbin domain, respectively (R, S). WDs are outlined by white dotted lines based on DAPI staining. Scale Bars: 100 μm. Data in charts represent means ± SD, \*\**p* < 0.01, \*\*\**p* < 0.001, \*\*\*\**p* < 0.0001, n.s. = non-significant. See also Supplementary Figure S3.

### The splicing defects caused by Ecd deficiency are Xrp1 independent

Intriguingly, despite the comprehensive rescue provided by Xrp1 or Irbp18 silencing, the absence of neither of the two TFs mitigated the lethality of animals with targeted Ecd knockdown in the nubbin domain. This suggests that Ecd plays an essential role distinct from governing the Xrp1-Irbp18 function and that Xrp1-Irbp18 upregulation is triggered as part of the adaptive response to damage inflicted by Ecd scarcity. Considering the involvement of Ecd in U5 snRNP and spliceosome assembly, we hypothesized that Ecd deficiency provokes changes to gene expression and splicing, independent of Xrp1-Irbp18 that are incompatible with successful development. To this end, we profiled and compared transcriptomes of control (*nub>*), *nub>ecd^RNAi^*, and *nub>ecd^RNAi^ xrp1^RNAi^*WDs using bulk mRNA-seq generated independently of that above (Supplementary Figure S4 and Supplementary Dataset S3). A highly significant overlap of DEGs in response to reduced Ecd between the two transcriptomic experiments (72%, *p* ≤ 0.0001, hypergeometric test) underscored the robustness of the molecular phenotype. Of the 2729 genes upregulated in response to Ecd deficiency compared to control (uniquely identified or overlapping genes, *pAdj* < 0.05), expression of 72% was significantly reduced upon simultaneous knockdown of *xrp1* in comparison to *ecd^RNAi^* alone (Figure 6A and Supplementary Dataset S3). The Xrp1 molecular footprint was even more pronounced among 2604 genes downregulated by *ecd* knockdown with 83% being re-expressed in *nub>ecd^RNAi^ xrp1^RNAi^* cells (Figure 6B). In agreement with the phenotypic rescue, and the role of Xrp1 in stress responses and in the maintenance of genome stability (Akdemir *et al*., 2007; Francis *et al*., 2016; Lee *et al*, 2018; Mallik *et al*, 2018), Xrp1-dependent DEGs showed enrichment for functions related to responses to DNA damage and repair, but also pre-mRNA splicing and RNA processing (Figure 6C, D and Supplementary Dataset S3). In contrast, the Xrp1-independent group of upregulated and downregulated genes revealed overrepresentation of genes implicated in the regulation of protein and enzyme activities, including proteasomal degradation (Figure 6C and Supplementary Dataset S3) and nucleobase metabolic and biosynthetic processes, respectively (Figure 6D and Supplementary Dataset S3).

**Figure 6.**
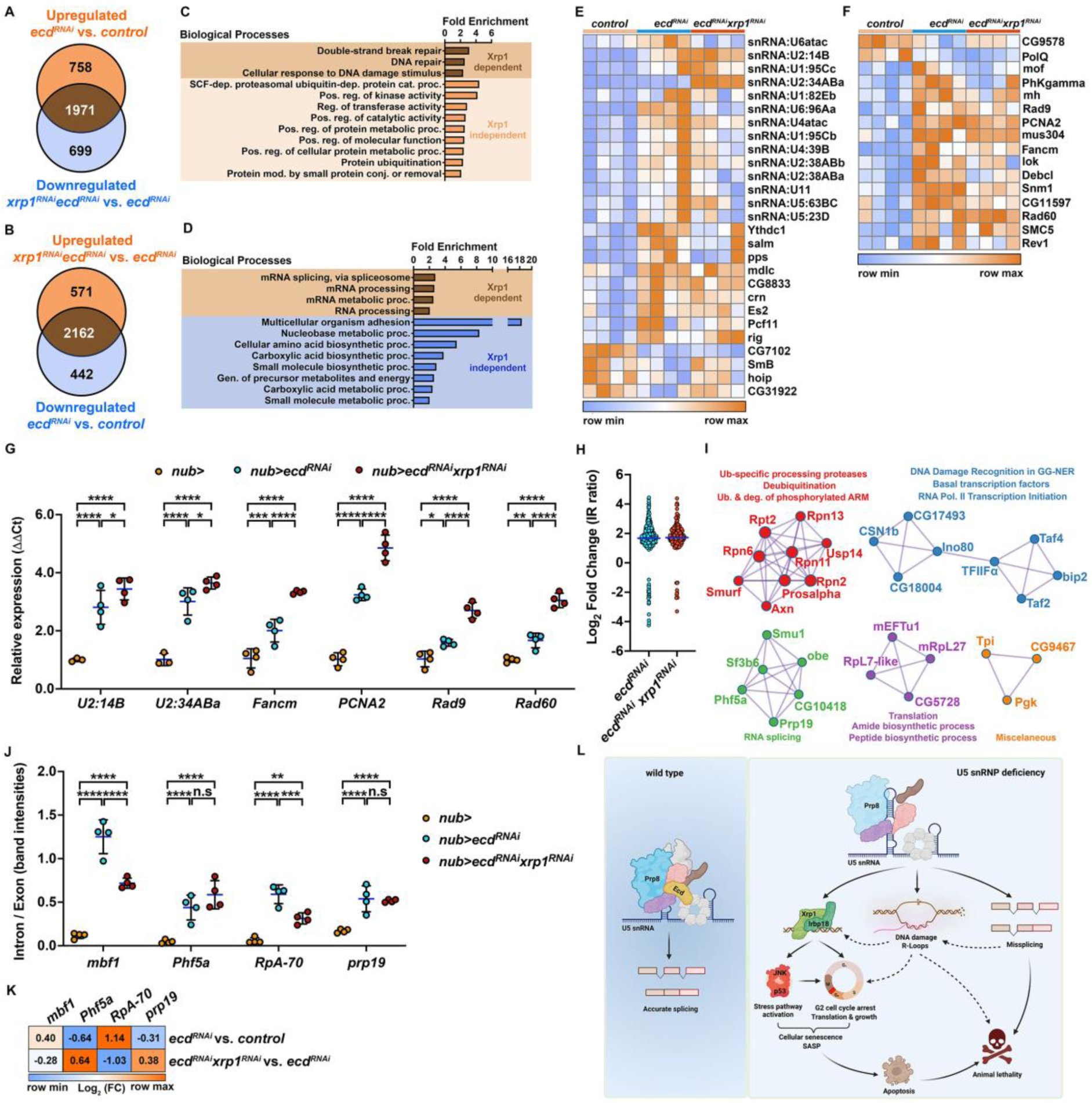
Splicing defects inflicted by Ecd deficiency are Xrp1-independent. **(A-B)** Venn diagrams show an overlap between genes upregulated (A) and downregulated (B) in Ecd deficient (*nub>mRFP ecd^RNAi^*) WDs compared to controls (*nub>mRFP*), and those downregulated (A) and upregulated (B) in double deficient (*nub>mRFP ecd^RNAi^ xrp1^RNAi^*) WDs compared to *ecd* knockdown alone. Genes at the intersection of the two groups are considered Xrp1-dependent. **(C-D)** GO enrichment plots for significantly upregulated (C) and downregulated (D) genes (*pAdj* < 0.05) in *nub>mRFP ecd^RNAi^* WDs compared to controls (*nub>mRFP*) whose expression is Xrp1-dependent and independent. Plots show fold enrichment of significant functional GO terms as identified by ShinyGO v0.76. Only GO categories with ≥ 5 genes and Fold Enrichment ≥ 2 are shown. Redundant GO terms were eliminated. **(E-F)** Expression of Xrp1-independent genes implicated in the cellular response to DNA damage stimulus (F) and pre-mRNA splicing (E) in the WDs samples of indicated genotypes. Heatmaps represents the Z-scores of normalized counts across rows. **(G)** Quantitative PCR of candidate transcripts involved in splicing and the DNA damage response, identified by the RNA-seq analysis as potentially Xrp1-independent. Knockdown of Xrp1 (*nub>mRFP ecd^RNAi^ xrp1^RNAi^*) did not restore the levels of the transcripts to control (*nub>mRFP*) levels. Data are fold changes in expression relative to control (ΔΔCt method). Levels of TATA binding protein (*tbp*) transcripts were used for normalization. Two-way ANOVA with Tukey’s multiple comparisons test was used to calculate *p*-values. Lines represent means ± SD. \**p* < 0.05, \*\**p* < 0.01, \*\*\**p* < 0.001, \*\*\*\**p* < 0.0001. **(H)** Fold changes (log_2_) of intron retention ratios significantly different (*pAdj* < 0.05) between control (*nub>mRFP*) and Ecd deficient (*nub>ecd^RNAi^*) or double deficient WDs (*nub>mRFP ecd^RNAi^ xrp1^RNAi^*) for each detected intron retention event. The blue line represents the mean. **(I)** Protein-protein interaction network enrichment analysis for genes misspliced in Ecd (Xrp1 independent) deficient cells. Neighborhoods of densely connected proteins as determined by the MCODE algorithm applied to the protein interaction networks generated by Metascape (Zhou *et al*., 2019). Functional enrichment analysis was applied to the MCODE networks (Bader & Hogue, 2003), and the three most significant GO terms were retained. **(J)** Semi-quantitative PCR on candidate genes shows increased intron retention in Ecd deficient WDs (*nub>mRFP ecd^RNAi^*) compared to the controls (*nub>mRFP*). The splicing defects persist upon *xrp1* knockdown (*nub>mRFP ecd^RNAi^ xrp1^RNAi^*). The intron/exon ratio was calculated based on intron- and exon-specific band intensities (see Supplementary Figure S5C). Two-way ANOVA with Tukey’s multiple comparisons test was used to calculate *p*-values. Data represent means ± SD. \*\**p* < 0.01, \*\*\**p* < 0.001, \*\*\*\**p* < 0.0001, n.s. = non-significant. **(K)** Fold changes (log_2_) of candidate genes with identified intron retention events by the IRFinder algorithm as compared to the control (*nub>mRFP*) WDs, or between the two experimental groups (*nub>ecd^RNAi^*and *nub>ecd^RNAi^ xrp1^RNAi^*). Knockdown of *xrp1* restored the expression of candidate genes deregulated in Ecd deficient WDs close to control levels. **(L)** Graphical summary. Intact U5 snRNP ensures constitutive and alternative splicing to generate transcript variants necessary for cell function, and normal animal development. Ecd deficiency interferes with U5 snRNP maturation causing aberrant splicing, gene dysregulation, DNA damage and R-loop accumulation, which trigger transcriptional upregulation of of Xrp1 and Irbp18. Xrp1-Irbp18 heterodimer acts as the key regulator of cellular stress signaling and the downstream responses, promoting senescence-like phenotype, including cell cycle arrest, increased translation rates, cellular growth and SASP. Chronic activation of Xrp1-Irbp18 ultimately leads to apoptosis and organismal death. In the absence of Xrp1-Irbp18-mediated response, loss of functional U5 snRNP remains lethal likely because of dysregulation and aberrant splicing of genes vital to the cellular homeostasis accompanied with accumulation of R-loops and DNA damage. See also Supplementary Datasets S3, S4 and Supplementary Figure S4, S5.

Although pre-mRNA splicing, RNA metabolism and DNA damage responses (DDR) emerged among the main processes controlled by Xrp1, a handful of splicing factors, spliceosome components and key regulators of genome stability remained dysregulated in *nub>ecd^RNAi^ xrp1^RNAi^* WDs (Figure 6E, F). RT-qPCR, using an independent set of biological samples, validated this Xrp1-independent upregulation, including the expression of the two *U2:14B* and *U2:34ABa* snRNA isoforms, *Fanconi anemia group M helicase* (*Fancm*), *Proliferating cell nuclear antigen 2* (*PCNA2*), *Rad60* and *Rad9* (Figure 6G and Supplementary Dataset S3 and S4). In fact, the expression of all six genes was further increased upon simultaneous knockdown of *xrp1* compared to *nub>ecd^RNAi^*WDs (Figure 6G). Importantly, a comparative splicing analysis using NxtIRFcore and IRFinder2 (Middleton *et al*, 2017), detected vast changes in pre-mRNA processing in both *nub>ecd^RNAi^*and *nub>ecd^RNAi^ xrp1^RNAi^* samples compared to control WDs (Supplementary Dataset S4). We found 413 differential splicing events (*pAdj* < 0.05) in *ecd^RNAi^* and 224 in *ecd^RNAi^ xrp1^RNAi^* samples, with many missplicing events being shared between the two genotypes. For example, 121 out of 212 introns and known exons retained in *nub>ecd^RNAi^* compared to control were also identified in *nub>ecd^RNAi^ xrp1^RNAi^* (Figure 6H and Supplementary Dataset S4). Strikingly, the GO enrichment analysis of the protein-protein interaction network generated from Xrp1-independent misspliced genes identified five MCODE networks comprised of components linked to RNA splicing, ubiquitin-mediated proteasomal degradation, DNA damage recognition, transcription, and translation (Figure 6I and Supplementary Dataset S4). The retention events were validated by the semi-quantitative RT-PCR on selected candidate genes involved in replication (*RpA-70*), regulation of stress-induced gene expression (*mbf1*), RNA splicing (*prp19, Phf5a*) and genome maintenance and transcription (*prp19*). Consistent with the genome-wide splicing analysis, all four candidates were misspliced in response to reduced Ecd (Figure 6J and Supplementary Figure S5A, B) and this aberrant pattern persisted upon simultaneous knockdown of *ecd* and *xrp1* (*nub>ecd^RNAi^ xrp1^RNAi^*), although for *mbf1* and *RpA-70* it was not to the same extent (Figure 6J). Importantly, the expression of the tested candidates was restored close to control levels by *xrp1* knockdown (Figure 6K and Supplementary Dataset S3), thus minimizing the potential confounding effect of differential gene expression on splicing. Collectively, these data provide evidence for a vital role of Ecd in controlling both the expression and pre-mRNA splicing of key sensors and regulators of genome, transcriptome, and proteome integrity, independent of Xrp1-Irbp18.

## DISCUSSION

The U5 snRNP forms the heart of the active spliceosome and is indispensable for splicing all introns in multicellular eukaryotes. A complete absence of individual U5 snRNP components causes early embryonic lethality (Bujakowska *et al*, 2009; Gaziova *et al*, 2004; Keightley *et al*, 2013; Kim *et al*., 2009) and loss of function clones cannot be recovered or are swiftly eliminated from mitotically active tissue (Andersen & Tapon, 2008; Claudius *et al*., 2014; Erkelenz *et al*., 2021; Gaziova *et al*., 2004). However, mutations in, and deregulation of, U5 snRNP components have been linked to highly tissue-specific pathologies as diverse as retinopathies, craniofacial disorders, and cancer (Adler *et al*, 2014; Krausová & Staněk, 2018; Kurtovic-Kozaric *et al*, 2015; Mirza *et al*, 2022; Wood *et al*., 2021; Xu *et al*, 2018; Zhao *et al*, 2012). Here, we used the developing wing imaginal epithelium of *Drosophila* to determine the cascade of events triggered by U5 snRNP scarcity caused either by targeted knockdown of the U5 snRNP biogenesis factor Ecd, or overexpression of an RP13-associated pathogenic variant of Prp8. Our results show that the lack of functional U5 snRNP, in the mitotically active cells of the nubbin domain, triggers a stress response program which engages the protective function of p53- and JNK-mediated signaling until their prolonged activity becomes toxic and induces apoptosis. Using a paradigm of Ecd deficiency, we establish Xrp1 and its dimerizing partner Irbp18 as the primary upstream inducers of p53 and JNK signaling, and mediators of the cellular responses to damage caused by U5 snRNP malfunction. In mammals, C/EBP transcription factors have been recognized as important regulators of genes associated with senescence and senescence-associated secretory phenotype (SASP) (Kumari & Jat, 2021). We provide functional genetic evidence that the increased expression of Xrp1-Irbp18 in Ecd deficient cells drives cell behaviors characteristic for cellular senescence and SASP (Cosolo et al., 2019; Kumari and Jat, 2021), including G2-phase cell cycle arrest, DNA damage, increased translation rate, cell growth, and paracrine secretion. Importantly, besides governing the canonical stress signaling and cellular properties of Ecd deficient cells, Xrp1-Irbp18 activation reduces proliferation also in neighboring imaginal territories, distorting organ proportions. We propose that this non-autonomous signaling could be mediated through a secretion of a systemic *Drosophila* insulin-like peptide 8 (Dilp8) (Boulan *et al*., 2019) whose expression was dramatically increased in Ecd deficient cells (Supplementary Dataset S2). Silencing of Xrp1-Irbp18 abolishes the stress-induced senescent phenotypes of Ecd deficient cells, revealing the pervasive missplicing of critical regulators of transcription, splicing, DNA damage, and proteostasis, which may in turn trigger Xrp1-Irbp18 upregulation and result in animal lethality (Figure 6L).

Underscoring a functional and physical coupling of transcription and splicing (Nojima *et al*, 2018; Saldi *et al*, 2016; Tellier *et al*., 2020), our molecular profiling of WDs comprised of Ecd deficient cells revealed a vast dysregulation of both transcription and splicing. Our analysis showed that changes in the *nub>ecd^RNAi^*transcriptome were dominated by aberrant expression of protein-coding genes. However, we also detected widespread de-repression of transposons, mostly residing in heterochromatic regions, indicating a possible disruption of chromatin organization and/or the mechanisms required for TE silencing. Interestingly, excessive TE activation has been observed in senescent human cells and in response to aging in both *Drosophila* and mammals (Colombo *et al*, 2018; De Cecco *et al*, 2013; Li *et al*, 2013; Rigal *et al*, 2022). Moreover, a recent analysis of RNA-seq datasets of senescent or aged cells in both humans and mice correlated increased transposon expression with transcriptional readthrough and intron retention (Pabis *et al*, 2023).

Importantly, the molecular signature of Ecd deficient WDs correlated with our observed phenotypes. The accumulation of DNA damage markers e.g., pH2Av was accompanied by the upregulation of several genes directly involved in DNA damage sensing and repair pathways. These sensing and repair pathways included homologous recombination (HR, *mre11, Rad50, nbs, RPA3*), non-homologous end joining (NHEJ, *Irbp*, *Ku80, Xrp1*, *Irbp18*, *DNAlig3*, *DNAlig4*, *RPA1*), global-genome nucleotide excision repair (GG-NER, *mrn*, *mus201 Xpc, Ercc1*) and Fanconi anemia pathway required for a resolution of highly toxic interstrand crosslinks (ICLs, *FANCI*, *FANCM*, *DNApol-iota*, *DNApol-zeta*). The increased translation rate was supported by upregulation of genes coding for small and large ribosomal subunits as well as regulators of ribosome biosynthesis and cytoplasmic translation. In contrast, genes associated with the spliceosome, proteasome function, protein processing, and signaling pathways controlling epithelial morphogenesis and differentiation were downregulated. The upregulation of autophagy-related genes could indicate a compensation for proteasome impairment (Dikic, 2017) and aligns with accelerated lysosome-dependent degradation of Prp8 in absence of Ecd (Erkelenz *et al*., 2021). Importantly, autophagy is among the effector mechanisms of senescence required for acquisition of the senescence phenotype (Young *et al*, 2009). Interestingly, p53 can promote autophagy (Kenzelmann Broz *et al*, 2013; Robin *et al*, 2019) and switches from a pro-apoptotic inducer to a pro-survival factor in G2-arrested cells (Ruiz-Losada *et al*, 2022). Together, this suggests that Xrp1-dependent p53 upregulation could exert a cytoprotective role in splicing deficient cells by inducing autophagy and delaying proteotoxicity caused by increased translation and proteasome impairment, promoting survival and maintenance of irreversibly damaged cells with their apoptotic removal.

Strikingly, >70% of genes differentially expressed in response to reduced Ecd levels were dependent on Xrp1, underscoring Xrp1 as the major driver of stress-induced senescent cellular behaviors. However, given the extensive deregulation of transcription factors in *nub>ecd^RNAi^* wing discs, we propose that the normalization of gene expression upon knockdown of *xrp1* results from both loss of regulation by Xrp1, and a rewiring of the TF network, including p53 and JNK-controlled transcriptional targets. Consistent with its role in protecting genome integrity (Akdemir *et al*., 2007; Brodsky *et al*., 2004; Francis *et al*., 2016), the upregulation of DDR and repair genes in response to U5 snRNP scarcity required Xrp1. In contrast, pre-mRNA splicing via spliceosome and RNA metabolism were highlighted among processes downregulated by Xrp1. These results indicate that Xrp1 induction in Ecd deficient cells may be two sided, promoting DDR to alleviate the primary insult to genome integrity while aggravating spliceosome dysfunction. It is plausible that the chronic shortage of properly processed transcripts and/or translation of aberrantly spliced mRNAs could enhance the stress load in Ecd deficient cells. Intriguingly, some of the DDR and repair genes (e.g., *Irbp18, Ku80, RpA-70, and His2Av*) and splicing factors (e.g., *Prp19, x16*) have been identified as interacting partners of the large and small Xrp1 protein isoforms (Mallik *et al*., 2018). This is important to consider when interpreting our results. While the downregulation of pH2Av foci upon *xrp1* silencing in Ecd deficient cells could signal alleviation of DNA damage, it is also plausible that the DNA damage persists and cannot be properly recognized because of the reduced expression and/or failed recruitment of DNA damage factors in the absence of Xrp1.

There is now a multitude of experimental evidence linking deregulated splicing to DNA damage and genomic instability with pre-mRNA splicing and genomic stability influencing each other through several different mechanism, including aberrant splicing of DDR factors or the accumulation of RNA/DNA duplexes known as R-loops (Wickramasinghe & Venkitaraman, 2016). R-loops may prove to be beneficial in fine-tuning gene expression, however, their ectopic formation and persistence due to spliceosome dysfunction are a significant source of DNA damage and genomic instability from budding yeast to mammals (Bonnet *et al*, 2017; Chakraborty *et al*, 2018; Halme *et al*, 2010; Li & Manley, 2005; Paulsen *et al*., 2009; Tresini *et al*., 2015). The increase in the number and strength of RNase H1 chromatin binding events identified by our TaDa profiling together with Xrp1-independnt upregulation of several genome caretakers required for R-loop removal, such as *Fanconi anemia group M helicase* (*Fancm*), *Rad60*, *Proliferating cell nuclear antigen 2* (*PCNA2*) and *Smc5* (Chang *et al*, 2019; Hodson *et al*, 2018; Kim *et al*, 2020; Lafuente-Barquero *et al*, 2017) supports the notion that R-loops may accumulate in response to Ecd loss. The contribution of R-loops to the phenotypes of Ecd deficient cells is a matter of future investigation. Importantly, splicing analyses highlighted Xrp1-independent missplicing of genes pivotal to transcription, splicing, DNA damage, and proteostasis. In this context, missplicing of *Prp19* and components of the chromatin remodeling Ino80 complex (Ino80C, *Ino80* and *CG18004* - Ino80 complex subunit E) are particularly intriguing. *Prp19* encodes an evolutionarily conserved splicing factor with E3 ubiquitin ligase activity. Prp19 was initially described as a part of the NineTeen Complex (NTC) required for the catalytic activation of the spliceosome (David et al., 2011). However, Prp19 also plays an essential role during transcriptional elongation and DNA repair by promoting recruitment and stabilization of the THO/TREX (TRanscription-EXport) complex and DNA repair factors (Chanarat & Sträßer, 2013). Ino80C is indispensable for nucleosome remodeling during transcription, replication, and DNA damage repair (Gurova *et al*, 2018; Poli *et al*, 2017). Proper function of THO/TREX and Ino80C have been shown to prevent the formation of harmful DNA-RNA hybrids (Domínguez-Sánchez *et al*, 2011; Huertas & Aguilera, 2003; Prendergast *et al*, 2020). Considering the pivotal role of Prp19 and Ino80C at the interface of DNA replication, transcription and RNA metabolism, their compromised function in Ecd deficient cells could contribute to molecular defects and organismal death prior to and independent of the activation of Xrp1-driven stress responses. Given that Prp19 has been detected within Xrp1 interactome (Mallik *et al*., 2018), it is tempting to speculate that Prp19 missplicing could affect Xrp1 levels and/or activity.

Our study contributes novel insights into the emerging role of Xrp1 as a stress-inducible transcription factor that governs molecular and cellular phenotypes in response to various stress insults, ranging from irradiation and P-element transposition (Akdemir *et al*., 2007; Brodsky *et al*., 2004; Francis *et al*., 2016), ER and proteotoxic stress (Baumgartner *et al*, 2021; Brown *et al*., 2021; Langton *et al*., 2021; Lee *et al*., 2018; Ochi *et al*, 2021; Recasens-Alvarez *et al*, 2021) to spliceosome malfunction (our study). Consistent with the published studies, our results show that Xrp1 levels must be tightly regulated as its overexpression or chronic activation is sufficient to induce cell death (Baillon *et al*, 2018; Baumgartner *et al*., 2021; Blanco *et al*., 2020; Boulan *et al*., 2019; Langton *et al*., 2021; Recasens-Alvarez *et al*., 2021). However, we also demonstrate that the cellular context, in which Xrp1 operates is essential for the phenotypic outcome. Xrp1 has been implicated in the neurotoxicity and motor defects in a *Drosophila* model of Amyotrophic lateral sclerosis (ALS) based on loss of an RNA binding protein Cabeza (Caz) or overexpression of its mutant human ortholog FUS (Mallik *et al*., 2018). More recently, Xrp1 was defined as a master regulator of Minute cell competition caused by ribosomal protein haploinsufficiency (“Minute” - *Rp*/+ cells) (Baillon *et al*., 2018; Boulan *et al*., 2019; Langton *et al*., 2021; Lee *et al*., 2018; Ochi *et al*., 2021).

In all cases, Xrp1 upregulation drives significant transcriptome remodeling and the Xrp1-controlled gene sets show marked overlap among Ecd-deficient and *Rp*/+ imaginal discs as well as *caz* mutant larval brains (Lee *et al*., 2018; Mallik *et al*., 2018 and our data). The phenotypic consequences of Xrp1 induction are, however, different, and sometimes opposing. In *Rp*/+ cells, Xrp1 upregulation reduced global translation rate and cell growth by engaging in proteotoxic stress-induced PERK-eIF2α signaling resulting in apoptotic elimination of cells with lower fitness. Reducing Xrp1 levels restored both translation and growth of *Rp*/+ cells allowing their survival surrounded by wild type neighbors. In contrast, in U5 snRNP deficient cells, Xrp1 triggers senescence program promoting translation and growth, while apoptosis is delayed until irreparable damage and breakdown of proteostasis override the pro-survival mechanisms. Supporting our findings, recent single cell profiling studies define Xrp1 as a shared molecular hallmark of senescence cell clusters in wounded and tumor bearing imaginal tissue (Floc’hlay *et al*, 2022). We propose that the differential impact of Xrp1 upregulation on protein synthesis and growth may lie in different stress signals to which Xrp1 initially responds and the cell cycle stage or cellular milieu in which it operates. Unlike Ecd deficient cells, Rp/+ cells might be prone to proteotoxicity due to aggregation of orphan ribosomal proteins (Baumgartner *et al*., 2021; Recasens-Alvarez *et al*., 2021) that will poise Xrp1 activity towards amplification of PERK-eIF2α signaling turning off translation. When surrounded by wild type cells, Rp/+ cells will be eliminated. Although experimental evidence pinpointed RpS12 protein to activate Xrp1 in Rp/+ cells (Boulan *et al*., 2019; Ji *et al*, 2019; Kale *et al*, 2018; Lee *et al*., 2018), the molecular mechanism remains unknown. Importantly, stimulating autophagy in RP-deficient cells proved beneficial reducing proteotoxic stress and tissue damage (Recasens-Alvarez *et al*., 2021). In cells experiencing U5 snRNP shortage, the trigger for Xrp1 upregulation remains to be determined. We suggest that the simultaneous activation of p53 and JNK signaling downstream of Xrp1 is crucial to the senescent phenotype of Ecd deficient cells.

Activation of p53 and JNK enforce G2 cell cycle arrest that is favorable to increased protein synthesis, growth, DNA damage repair and proteostasis network rewiring while reduces sensitivity to apoptosis (Cosolo *et al*., 2019; Ruiz-Losada *et al*., 2022). We envision that aggravation of proteotoxic stress due to chronic Xrp1 activity and/or lifting the cell cycle break would drive cell death.

In summary, our study demonstrates the requirement for the functional spliceosome in the maintenance of cellular, tissue, and organismal homeostasis. We identify Xpr1 as a key driver of stress responses induced by U5 snRNP scarcity and implicate cellular senescence controlled by the C/EBP family bZIP transcription factors in pathologies caused by U5 snRNP malfunction. Given the emerging evidence for pre-mRNA splicing as an essential contributor to the aging process, splicing factor dysregulation observed during aging could contribute to senescent cell accumulation, promoting the decline of cell and tissue functions and the development of age-related disorders.

## MATERIALS AND METHODS

### Fly stocks and husbandry

*Drosophila* stocks and crosses were maintained at 25 °C (unless specified otherwise) on a diet consisting of 0.8% agar, 8% cornmeal, 1% soymeal, 1.8% dry yeast, 8% malt extract and 2.2% sugar-beet syrup, supplemented with 0.625% propionic acid and 0.15% Nipagin. The following fly stocks were used: *w^1118^* (BDSC; RRID:BDSC_3605), *ecd^1^* (Garen *et al*, 1977), *prp8^del14^*(Stankovic *et al*., 2020), *UAS-Prp8^[S2178F]^* (Stankovic *et al*., 2020), *nub-Gal4, UAS-myr-mRFP* (*nub>mRFP*) (Erkelenz *et al*., 2021), *nub-Gal4* (Erkelenz *et al*., 2021), *UAS-ecd^RNAi^* (Claudius *et al*., 2014), *UAS-xrp1^RNAi^ #1* (BDSC; RRID:BDSC_34521), *UAS-xrp1^RNAi^ #2* (VDRC, ID 107860), *UAS-Irbp18^RNAi^*(VDRC ID 101871), *UAS-p35* (BDSC; RRID: BDSC_5072), *tub-Gal80^TS^* (BDSC; RRID:BDSC_7017), *UAS-LT3-Dam* (Southall *et al*., 2013), *UAS-LT3-Dam-RNase H1^WT^*(this study, see below), *ubi-FUCCI* (BDSC; RRID:BDSC_55124), *TRE-dsRED-2R* (Chatterjee & Bohmann, 2012), *xrp1-lacZ* (BDSC; RRID: RRID:BDSC_11569), *gstD1-GFP* (Sykiotis & Bohmann, 2008), *p53::GFP^[FlyFos]^*(VDRC, ID 318453), *egr::GFP^[FlyFos]^* (VDRC, ID 318615), *nos-phiC31, attP2* (BDSC; RRID:BDSC_25710), *Irbp18::GFP^[FlyFos]^* (VDRC, ID 318564), *Irbp::GFP^[FlyFos]^* (VDRC, ID 318033). See Supplementary Table S1 for all stock and recombinant line genotypes used in this study.

### Tissue dissection and immunostaining

Wing imaginal discs (WDs) were dissected on day 7 AEL (unless specified otherwise) in PBS and fixed using 4% paraformaldehyde in PBS containing 0.1% TritonX-100 (PBS-T) for 25 minutes at room temperature. Blocking was performed with 0.3% BSA in PBS-T for 30 minutes at room temperature. Incubation with primary antibodies diluted in the blocking solution was performed overnight at 4 °C while nutating. The following primary antibodies were used in this study: anti-RFP (rabbit, 1:1000;

MBL #PM005; RRID:AB_591279), anti-GFP (rabbit, 1:500; Thermo Fisher Scientific #G10362; RRID:AB_2536526), anti-GFP (goat, 1:500; Abcam #ab6673; RRID:AB_305643), anti-p53 antibody mix (mouse, 1:200; DSHB #p53 25F4; RRID:AB_579787, DSHB #p53 7A4, RRID:AB_579786), anti-phospho-Histone H2AvD (rabbit, 1:300; Rockland #600-401-914S; RRID:AB_11183655), anti-Ecd C-terminal (rat, 1:700) (Claudius *et al*., 2014), anti-Ecd N-terminal (rat, 1:700) (Claudius *et al*., 2014), anti-Fibrillarin (rabbit, 1:200; Abcam #ab5821; RRID:AB_2105785), anti-Lamin (mouse, 1:1000; DSHB #adl67.10; RRID:AB_528336), anti-phospho-histone H3 (rabbit, 1:500; Cell Signaling Technology #9701; RRID:AB_331535), anti-Dcp-1 (rabbit, 1:500; Cell Signaling Technology #9578; RRID:AB_2721060), anti-Mmp1 antibody mix (mouse, 1:100; DSHB #14A3D2, #3A6B4, #3B8D12; RRID:AB_579782, RRID:AB_579780, RRID:AB_579781), anti-puromycin (mouse, 1:1000; Millipore #MABE343, RRID:AB_2566826), anti-beta-Galactosidase (mouse, 1:100; DSHB #jie7, RRID:AB_528101). After five washes in PBS-T, staining with the appropriate secondary antibodies, diluted 1:1000 in blocking solution, was performed overnight at 4 °C while nutating: donkey anti-rabbit Alexa Fluor 488 (Jackson ImmunoResearch #111-545-144; RRID:AB_2338052), donkey anti-goat Cy2 (Jackson ImmunoResearch #705-225-147; RRID:AB_2307341), donkey anti-rabbit Cy5 (Jackson ImmunoResearch #711-175-152; RRID:AB_2340607), donkey anti-mouse Cy5 (Jackson ImmunoResearch #715-175-151; RRID:AB_2340820), donkey anti-rat Cy5 (Jackson ImmunoResearch #712-175-153; RRID:AB_2340672), donkey anti-mouse Cy3 (Jackson ImmunoResearch #715-165-151; RRID:AB_2315777), donkey anti-rat FITC (Jackson ImmunoResearch #712-095-153; RRID:AB_2340652).

### Image acquisition and processing

Images of *Drosophila* WDs were acquired with the Olympus FV-1000 confocal microscope using 20x UPlan S-Apo (NA 0.85), 40x UPlanApo (NA 1.30) and 60x UPlanApo (NA 1.35) objectives. All micrographs shown are maximum Z-projections generated with FluoView FV-10ASW software using 1.2 or 1.4 um step size unless specified otherwise (Olympus, RRID:SCR_014215). Panel assembly and brightness/contrast adjustments were done using Adobe Photoshop CC (Adobe Systems Inc., RRID:SCR_014199). Images used for quantification of fluorescence intensity were acquired using the same settings. WD size was measured by manual selection of the entire WD or the nubbin domain marked by mRFP and expressed as μm^2^. Quantification of pH3 foci was performed on maximum projections of confocal Z-stacks using the “Analyze Particles” command within ImageJ FIJI with a minimum size cutoff of 3 μm^2^ after image thresholding using the maximum entropy method (Kapur *et al*, 1985). Apoptosis was measured as the total area of Dcp-1 signal within the nubbin domain in maximum projections of confocal Z-stacks, after thresholding (default method). Levels of endogenous p53 or GFP-tagged p53 are represented as ratios of mean gray values of the fluorescence within and outside of the nubbin domain. Nucleolar sizes were determined based on immunostaining with a Fibrillarin antibody. Images were first de-noised (“Remove speckles” function), Gaussian blur was applied (Radius value of 2) and background was subtracted (Rolling ball radius of 20 pixels). Images were thresholded and the area of nucleoli was measured. Cell sizes were measured by manual selection of the membrane mRFP signal driven by the *nub-Gal4* driver. For each biological replicate, all cells spanning the diagonal corners of a micrograph were measured. All quantification related processing and measurements were performed with ImageJ FIJI (RRID:SCR_002285).

### Puromycin incorporation assay and quantification

Third instar wandering larvae (day 7 AEL) were inverted in PBS, and the intestine and fat body were removed. The tissues were transferred to a solution of 5 ug/mL of puromycin (Thermo Fisher Scientific #A1113803) in Shield and Sang M3 insect media (Sigma-Aldrich #S8398) containing 10% FBS (Gibco, Life Technologies) for 20 minutes. Samples were then fixed in 4% PFA in PBS-T for 40 minutes and processed for immunostaining as described above. Regions of interests were outlined based on the *nub>mRFP* signal and the mean gray values of puromycin antibody signal measured. The mean gray values of the hinge and notum regions of the WDs were used to calculate the fluorescence signal intensity ratio.

### EdU incorporation assay

The EdU incorporation assay was performed using the Click-iT Plus EdU cell proliferation kit for imaging (ThermoFisher Scientific #C10638). Third instar wandering larvae (day 7 AEL) were inverted in PBS and incubated in 15 μM EdU in PBS for 20 minutes at room temperature with gentle shaking. Upon washing twice with 3% BSA in PBS, samples were fixed in 3,7% formaldehyde for 15 minutes at room temperature. Samples were then washed and permeabilized using PBS-T. After washing twice with 3% BSA in PBS, the Click-It reaction cocktail was added to the samples, according to manufacturer’s recommendations (30 min) at room temperature protected from light. The samples were washed once, mounted, and imaged as described above.

### RNA isolation and RNA sequencing

150 WDs were dissected from third instar larvae (day 7 AEL) in PBS and snap-frozen in liquid nitrogen. Total RNA was isolated according to standard TRI Reagent protocol (Sigma Aldrich, #T9424), followed by DNase I treatment (Invitrogen, #AM2238) and repurification as described in (Mundorf & Uhlirova, 2016). For RNA sequencing, 2 μg of total RNA was used for library preparation (Illumina TruSeq Stranded total RNA Ribo-Zero) with four biological replicates per experimental group. Pair-end sequencing (100 bp) was performed using the Illumina HiSeq 2000 platform. Image analysis and base calling were carried out with the Illumina RTA software at run time.

### Differential gene expression and splicing analyses

Initial quality control of the raw data was performed using *FastQC* (RRID:SCR_014583). Illumina sequencing adapters were removed from the sequencing reads with the *cutadapt* tool version 3.5 (Martin, 2011) (RRID:SCR_011841). For differential gene expression analysis, *kallisto* v0.46.1 (Bray *et al*, 2016) (RRID:SCR_016582) was used for pseudoalignment of obtained reads to the complete *Drosophila melanogaster* transcriptome (BDGP6.28, Ensembl release 102) and transcript-level abundance quantification using 100 bootstrap samples. Summarization of the obtained estimated counts to gene-level was performed using *tximport* v1.22.0 (Soneson *et al*, 2015) (RRID:SCR_016752). Differential gene expression was determined using *DESeq2* v1.34.0 (Love *et al*, 2014) (RRID:SCR_015687). Heatmaps comparing expression levels of selected genes between samples were constructed from DESeq2 normalized counts and log2FC values using Morpheus (https://software.broadinstitute.org/morpheus/). Isoform-specific expression was assessed by *sleuth* v0.30.0 (Pimentel *et al*, 2017). For differential splicing analysis, the reads were aligned to the *Drosophila* melanogaster genome (BDGP6.28, Ensembl release 102) using *STAR* v2.7.9a (Dobin *et al*, 2013) (RRID:SCR_004463) in basic two-pass mode. The resulting alignments were loaded into NxtIRFcore *v1.2.1* for intron retention analysis on individual biological replicates in R. The resulting output tables were then used with the IRFinder *v2.0.0* (Middleton *et al*., 2017) DESeq2 wrapper for differential intron retention analyses using default settings.

Gene Ontology term enrichment analysis was performed using ShinyGO *v0.76* (Ge *et al*, 2020) using a custom background gene list containing all genes detected by the RNA-seq analysis. Fisher’s exact test with false discovery rate correction was used. Metascape *v3.5* (Zhou *et al*, 2019) was used to generate protein-protein interaction and MCODE networks (Bader & Hogue, 2003) using the default settings.

### Quantitative and semi-quantitative PCR intron retention assay

cDNA was synthesized from 2 ug of total RNA using GoScript Reverse Transcriptase (Promega, #A5003) with random hexamers according to the manufacturer’s protocol. Quantitative PCR was performed in three technical replicates and at least three biological replicates. Expression values were normalized to the levels of *tbp* transcripts (FlyBase ID: FBtr0071679). For calculation of the fold changes, ΔΔCT method was used (Livak & Schmittgen, 2001). Statistical significance of gene expression changes was determined using two-way ANOVA with Sidak’s correction for multiple comparisons.

For semi-quantitative PCR, a set of biological samples independent from those used for RNA sequencing was generated. cDNA was synthesized using the same protocol. Fragments of interest were amplified using the EmeraldAmp GT PCR Master Mix (Takara, #RR310B) and intron- or exon - specific forward primers combined with a common exon junction-specific reverse primer. The PCR products were resolved on a 3% agarose gels and stained with Ethidium bromide. Intensity profiles of the resulting bands were plotted using Image FIJI, and the area under the profile curves was used to calculate the intron/exon ratios. See Supplementary Table S2 for primer sequences.

### *In vivo* Targeted DamID

The coding sequence of *RNase H1* (isoform RA) was amplified from cDNA and cloned into the *pUASt-attB-LT3-NDam* vector (Southall *et al*., 2013) using the NotI and XhoI restriction enzymes (NEB #R0146S and #R0189S respectively). Transgenic flies were obtained by insertion of the *pUASt-attB-LT3-NDam-RNase H1* construct into the attP2 site on the third chromosome (68A4) using PhiC31 integrase-mediated transgenesis (BDSC; RRID:BDSC_25710). The expression of the UAS-based Dam-fusion constructs in the wing pouch using the *nub-Gal4* driver was induced by shifting the developing larvae from room temperature to 29 °C for 24 hours. This resulted in the inactivation of the temperature sensitive Gal80^TS^ while simultaneously rendering the mutant Ecd protein produced from the homozygous *ecd^1^* loci non-functional (Erkelenz *et al*., 2021). Third instar larvae were dissected on day 7 AEL. Samples were collected in four biological replicates, each containing 120 WDs. Sample processing was performed according to (Vogel *et al*, 2007) with modifications described in (Mundorf *et al*, 2019). Preparation for sequencing was performed as previously described (Mundorf *et al*., 2019). Sequencing libraries were generated using the Illumina TruSeq DNA kit library preparation protocol and sequenced on an Illumina HiSeq 2000 machine as pair-end 50 bp reads. The resulting sequencing reads files were trimmed using the *cutadapt* tool version 3.5 (Martin, 2011) (RRID:SCR_011841), and processed using the *damidseq_pipeline* v1.4.6 (Marshall & Brand, 2015) to obtain GFF files containing the normalized log_2_ ratio (Dam-fusion/Dam) for each GATC fragment in the *Drosophila* genome. Dam-RNase H1^WT^ binding was assessed using a peak finding algorithm based on the one described by Wolfram et. al. (2012). FDR was calculated for peaks (formed of two or more consecutive GATC fragments) for the individual replicates. Peaks with FDR < 0.01 were classified as significant. Additionally, a mean log_2_ ratio threshold of 0.5 for each peak was implemented for peak to gene association. Significant peaks present in all four replicates were used to form a final peak file. Any gene within 5 kb of a peak (with no other genes in between) was identified as a peak-associated gene. Read alignment files were converted into bedGraph format for coverage visualization using the *bamCoverage* tool with a bin size of 10 bp. Paired reads were extended, and data were normalized by 1x depth (reads per genome coverage, RPGC) with effective genome size set to 142573017 bp.

Averages of Dam-Fusion/Dam ratios for each GATC fragment from four biological replicates were used to generate the corresponding genome browser tracks. To identify differential binding events in each condition the difference between binding intensities for each genomic GATC fragment (represented as log_2_ Dam-RNase H1/Dam-only ratios) was calculated. Significant peaks with higher intensities in each condition compared to the other were then associated to the nearest gene (maximum distance of 5 kb).

## DATA AVAILABILITY

The raw and processed next generation sequencing data have been deposited in NCBI’s Gene Expression Omnibus (RRID:SCR 007303) (Edgar et al., 2002) and are accessible through GEO Series accession numbers GSE213325 for the Targeted DamID data (https://www.ncbi.nlm.nih.gov/geo/query/acc.cgi?acc=GSE213325), GSE203127 and GSE213298 for the RNA-Seq data (https://www.ncbi.nlm.nih.gov/geo/query/acc.cgi?acc=GSE213298), (https://www.ncbi.nlm.nih.gov/geo/query/acc.cgi?acc=GSE203127).

## ACKNOWLEDGEMENTS

We thank Alexandra Trifunovic Laboratory, the Bloomington *Drosophila* Stock Center supported by NIH grant NIH P40OD018537 (BDSC, Bloomington, IN, USA), the *Drosophila* Genomics Resource Center supported by NIH grant 2P40OD010949 (DGRC, Bloomington, IN, USA), the Zurich ORFeome Project (FlyORF, Zurich, Switzerland), the Vienna *Drosophila* Resource Center (VDRC, Vienna, Austria), and the Developmental Studies Hybridoma Bank (DSHB, Iowa City, IA, USA) for fly stocks, plasmids, and antibodies. We are grateful to Tina Bresser, Agnieszka Sokol, and Nils Teuscher for generation of transgenic lines, fly stock maintenance and technical assistance. This work was funded by UH 243/6-1 project to M.U and under Germany’s Excellence Strategy – CECAD, EXC 2030 – 390661388 both from the Deutsche Forschungsgemeinschaft (DFG, German Research Foundation).

## AUTHORS CONTRIBUTIONS

D.S. and M.U. conceived the study. D.S., L.T., and M.U. carried out the investigation. D.S., L.T., and M.U. performed formal analysis. D.S. wrote the initial draft. D.S., L.T., and M.U. wrote the manuscript and generated the visualizations therein. M.U. provided supervision, project administration, and funding.

## COMPETING INTERESTS

The authors declare no competing interests.

## SUPPLEMENTARY FIGURES

**Supplementary Figure S1.**
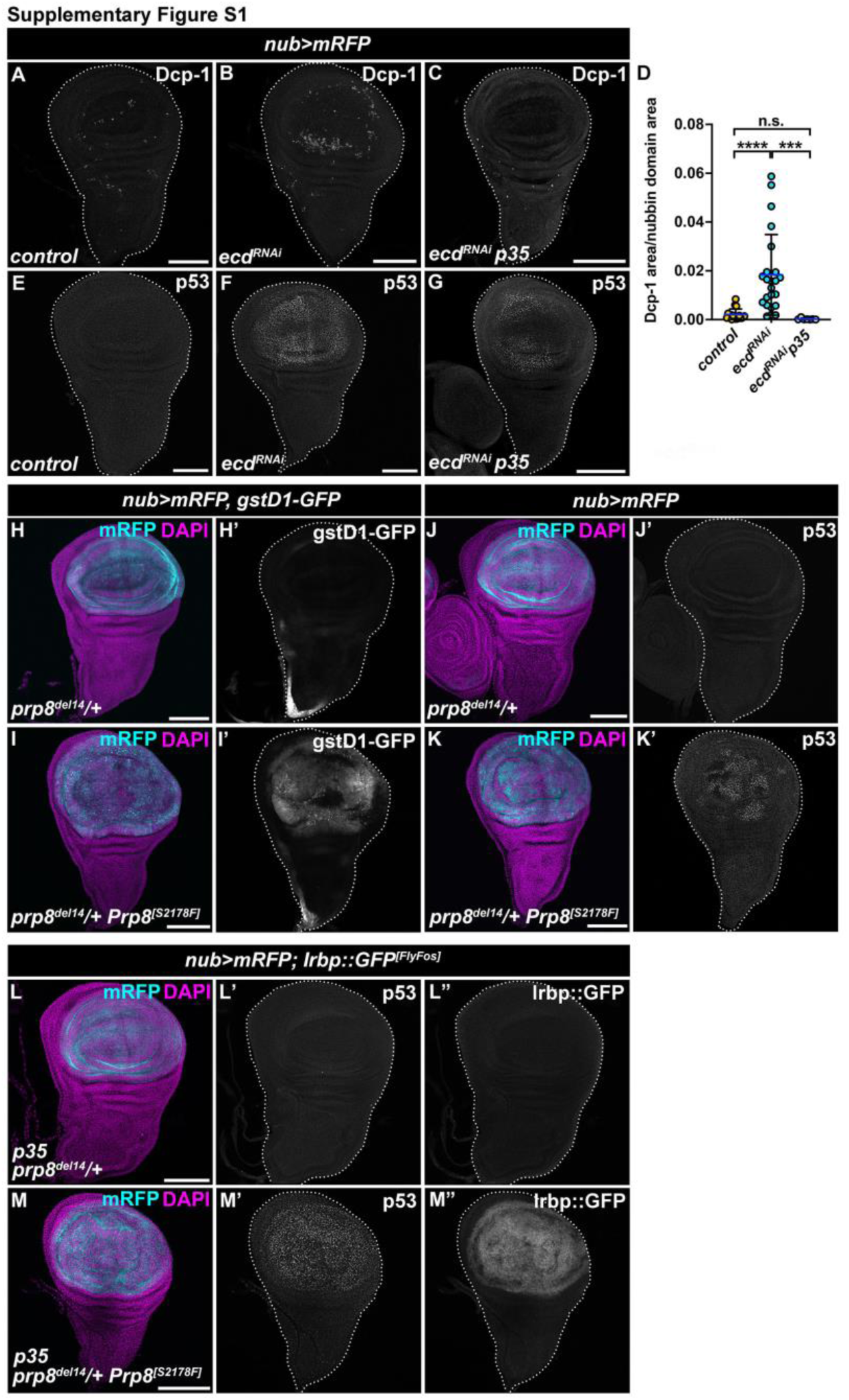
A pathogenic adRP Prp8 mutation recapitulates the phenotypes of Ecd deficiency. **(A-D)** Compared to control (*nub>mRFP*) (A), the wing pouch-specific knockdown of *ecd* (*nub>mRFP, ecd^RNAi^*) increases the levels of the effector caspase Dcp-1 (B) that is abolished by the overexpression of the pan-caspase inhibitor p35 (C). Quantification of Dcp1 positive area normalized to the wing pouch area in the control (*nub>mRFP*, n=18), *nub>mRFP ecd^RNAi^* (n=23) and *nub>mRFP ecd^RNAi^ p35* (n=7) WDs on day 7 AEL (D). One-way ANOVA with Tukey’s multiple comparisons test was used to calculate *p*-values. Data represent means ± SD, \*\*\**p* < 0.001, \*\*\*\**p* < 0.0001, n.s. = non-significant. **(E-G)** Compared to control (*nub>mRFP*) (E), the wing pouch-specific knockdown of *ecd* (*nub>mRFP ecd^RNAi^*) increases the levels and the endogenous p53 (F) that persist upon co-expression of a pan-caspase inhibitor p35 (G). **(H-K)** Compared to control (*prp8^del14/+^, nub>mRFP*) (H, J), the wing pouch-specific overexpression of *Drosophila* Prp8^[S2178F]^ (*prp8^del14/+^, nub>mRFP Prp8^[S2178F]^*) corresponding to human adRP-causing PRPF8^[S2118F]^ mutation induces the *gstD1-GFP* reporter construct (I’) as well as accumulation of endogenous p53 (K’). (L-M) The upregulation of endogenous p53 (M’) and the Irbp::GFP reporter construct (M’’) in wing pouch of *prp8^+/-^* heterozygous WDs overexpressing Prp8^[S2178F]^ persist after overexpression of p35. *Prp8^+/-^* heterozygosity itself does not induce p53 nor Irbp::GFP (L’, L’’). The micrographs are representative maximum intensity projections of multiple confocal sections of WDs on day 7 AEL. Nubbin domain and nuclei are marked by mRFP and DAPI, respectively. WDs are outlined by white dotted lines based on DAPI staining. Scale Bars: 100 μm.

**Supplementary Figure S2.**
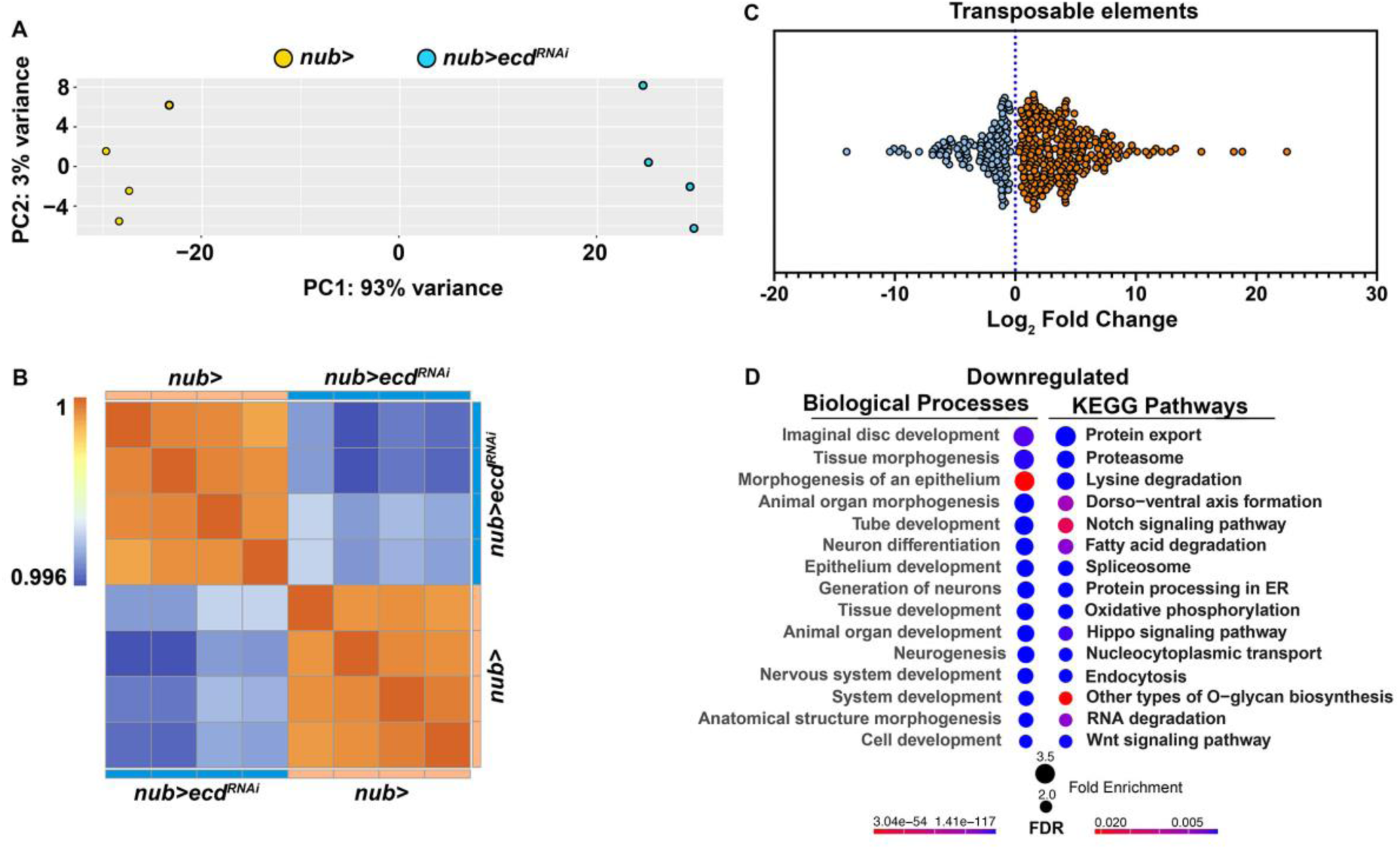
Ecd deficiency alters expression of transposable elements and downregulates genes involved in diverse developmental processes. **(A, B)** Principal component analysis (PCA) (A) and correlation plot (B) of all biological replicates, control (*nub>mRFP*) and Ecd deficient WDs (*nub>mRFP ecd^RNAi^*) used in the differential gene expression analysis shown in Figure 4. **(C)** Knockdown of *ecd* increases expression of transposable elements. Plot shows Log_2_ fold changes of significantly up- and down-regulated transposable elements in *nub>mRFP ecd^RNAi^* compared to control WDs (*nub>mRFP*). **(D)** GO enrichment plots for significantly downregulated genes (*pAdj* < 0.05) in Ecd deficient WDs (*nub>mRFP ecd^RNAi^*) compared to control (*nub>mRFP*) as identified by ShinyGO v0.76. as identified by ShinyGO v0.76. Only GO categories with ≥ 5 genes and Fold Enrichment ≥ 2 are shown. Redundant GO terms were eliminated. Related to Figure 4. See also Supplementary Dataset S2.

**Supplementary Figure S3.**
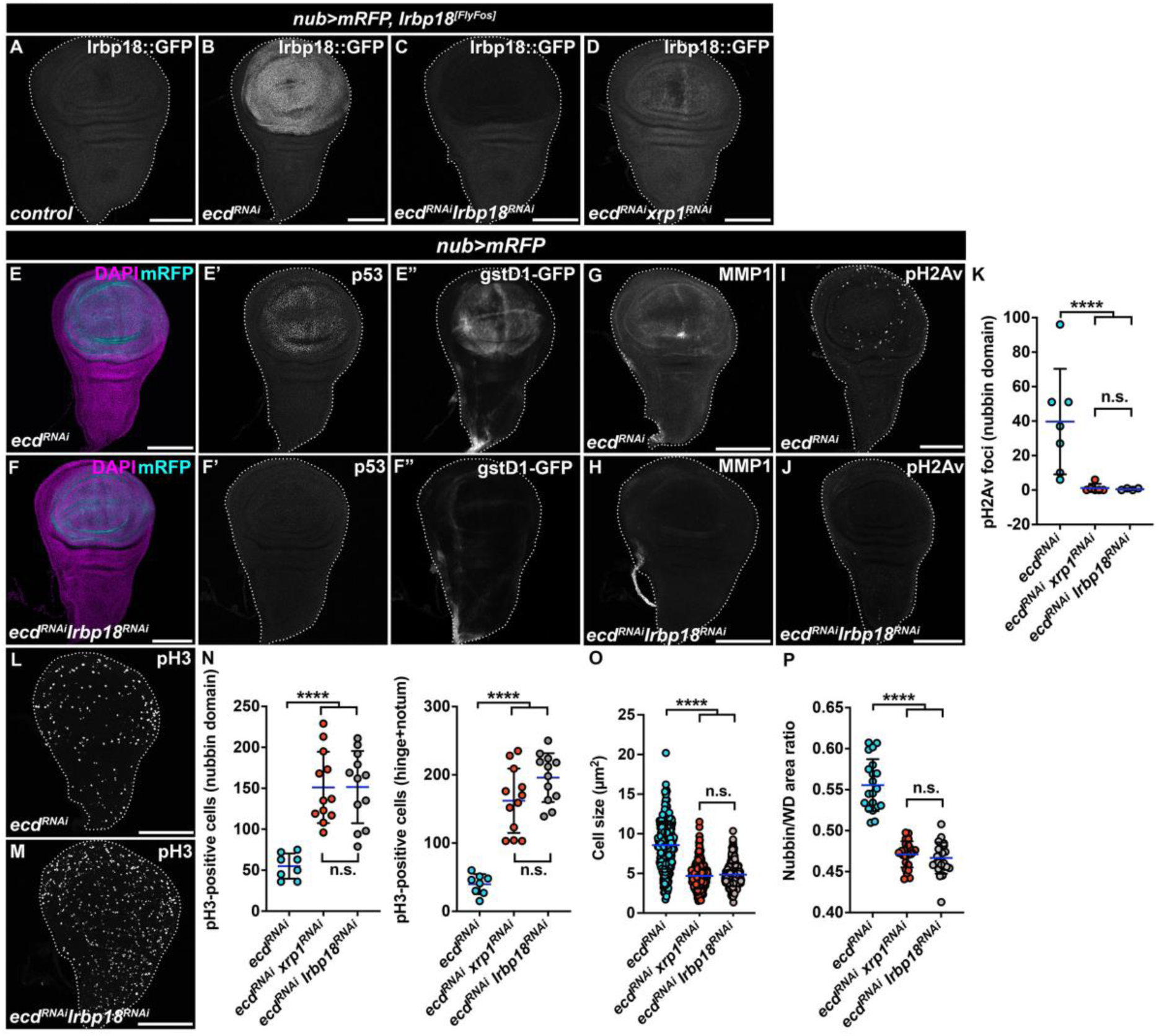
Stress response phenotypes induced by Ecd deficiency require Irbp18. **(A-D)** The upregulation of Irbp18 FlyFos reporter in Ecd deficient WD cells (*nub>mRFP ecd^RNAi^*) (B) compared to control (*nub>mRFP*) (A) is suppressed by simultaneous knockdown of *Irbp18* (*nub>mRFP ecd^RNAi^ Irbp18^RNAi^*) (C) or *xrp1* (*nub>mRFP ecd^RNAi^ xrp1^RNAi^*) (D). **(E-P)** Irbp18 silencing in Ecd deficient cells (*nub>mRFP ecd^RNAi^ Irbp18^RNAi^*) prevents the upregulation of p53 (F’), the gstD1-GFP reporter (F’’) and MMP1 (H) as observed in wing pouch cells of *nub>mRFP ecd^RNAi^* WDs (E’, E’’ and G, respectively). *Irbp18* knockdown also abolished the pH2Av fluorescent signal (J, G) present in the wing pouch of Ecd deficient WDs (I, G). Knockdown of Irbp18 restores cell proliferation within nubbin as well as hinge and notum region of *nub>ecd^RNAi^* WDs (L, M, N), as well as the cell size (O) and the size of the *nubbin* domain (P) recapitulating the effect of *xrp1^RNAi^*. pH3-positive cells were counted in the nubbin domain and hinge + notum regions in *nub>mRFP ecd^RNAi^* (n = 8), *nub>mRFP ecd^RNAi^ xrp1^RNAi^* (n = 12), and *nub>mRFP ecd^RNAi^ Irbp18^RNAi^* (n = 12) WDs. Sizes of cells spanning both diagonals of the wing pouch were quantified from the close-up confocal images (K, n=7 WD per genotype). Areas of the nubbin domain and the entire WDs were measured based on the mRFP and DAPI signals, respectively (L, n = 20-25). One-way ANOVA with Tukey’s multiple comparisons test was used to calculate *p*-values. Data represent means ± SD, \*\*\*\**p* < 0.0001, n.s. = non-significant. The micrographs are maximum intensity projections of multiple confocal sections of WDs at day 7 AEL. Nubbin domain and nuclei are marked by mRFP and DAPI, respectively (E, F). WDs are outlined by white dotted lines based on DAPI staining. Scale Bars: 100 μm. Related to Figure 5.

**Supplementary Figure S4.**
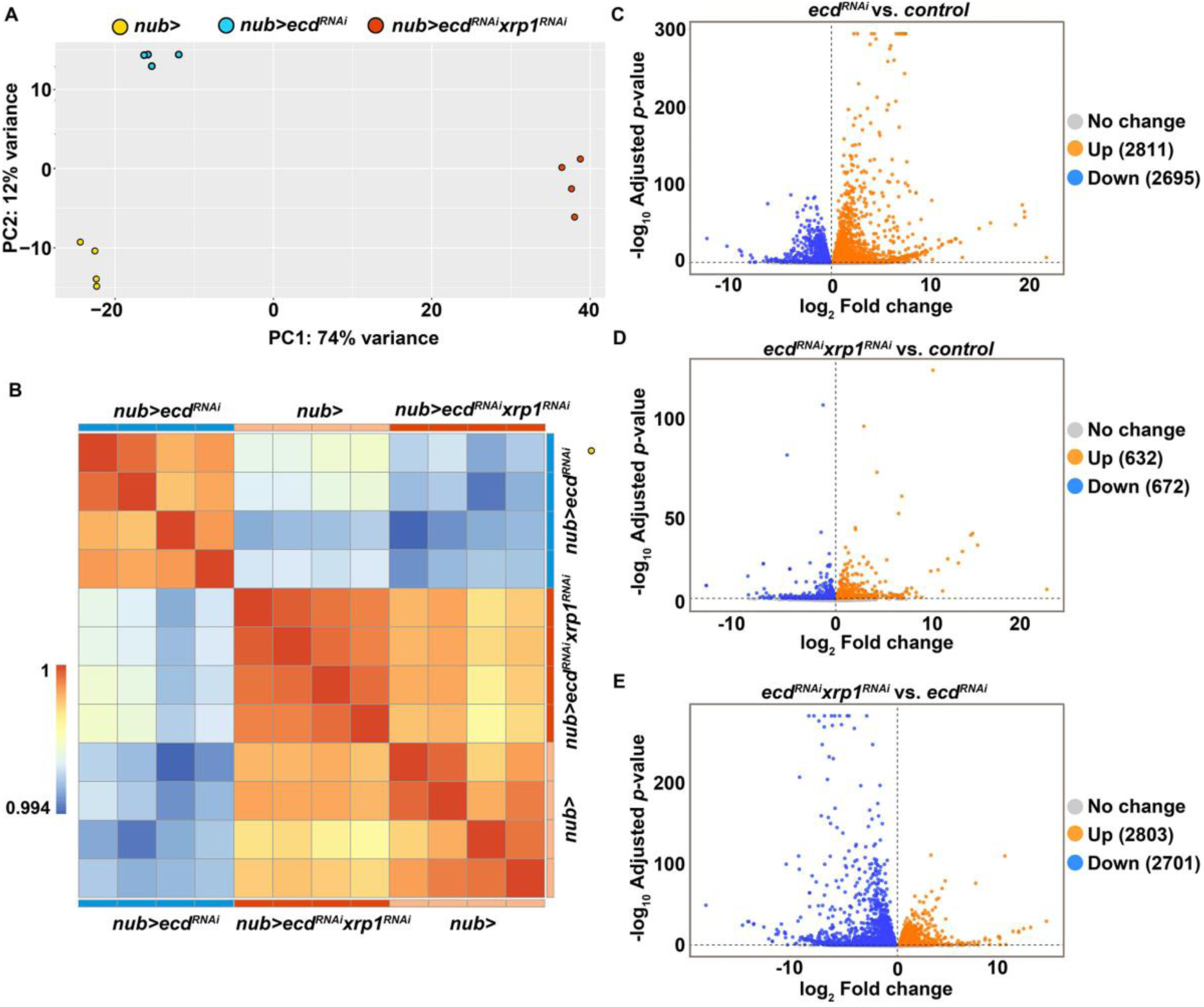
Gene expression changes in response to Ecd and Xrp1 deficiency. **(A, B)** PCA (A) and correlation plot (B) of all biological replicates from the three experimental genotypes, control (*nub>mRFP*), *nub>mRFP ecd^RNAi^* and *nub>mRFP ecd^RNAi^ xrp1^RNAi^*, used in the differential gene expression analysis shown in Figure 6. **(C-E)** Volcano plots depict all significantly regulated genes (*pAdj* < 0.05) in *nub>mRFP ecd^RNAi^* WD samples compared to controls (*nub>mRFP*) (C), *nub>mRFP ecd^RNAi^ xrp1^RNAi^* compared to controls (D) and *nub>mRFP ecd^RNAi^ xrp1^RNAi^* compared to *nub>mRFP ecd^RNAi^* WDs (E). Related to Figure 6. See also Supplementary Dataset S3.

**Supplementary Figure S5.**
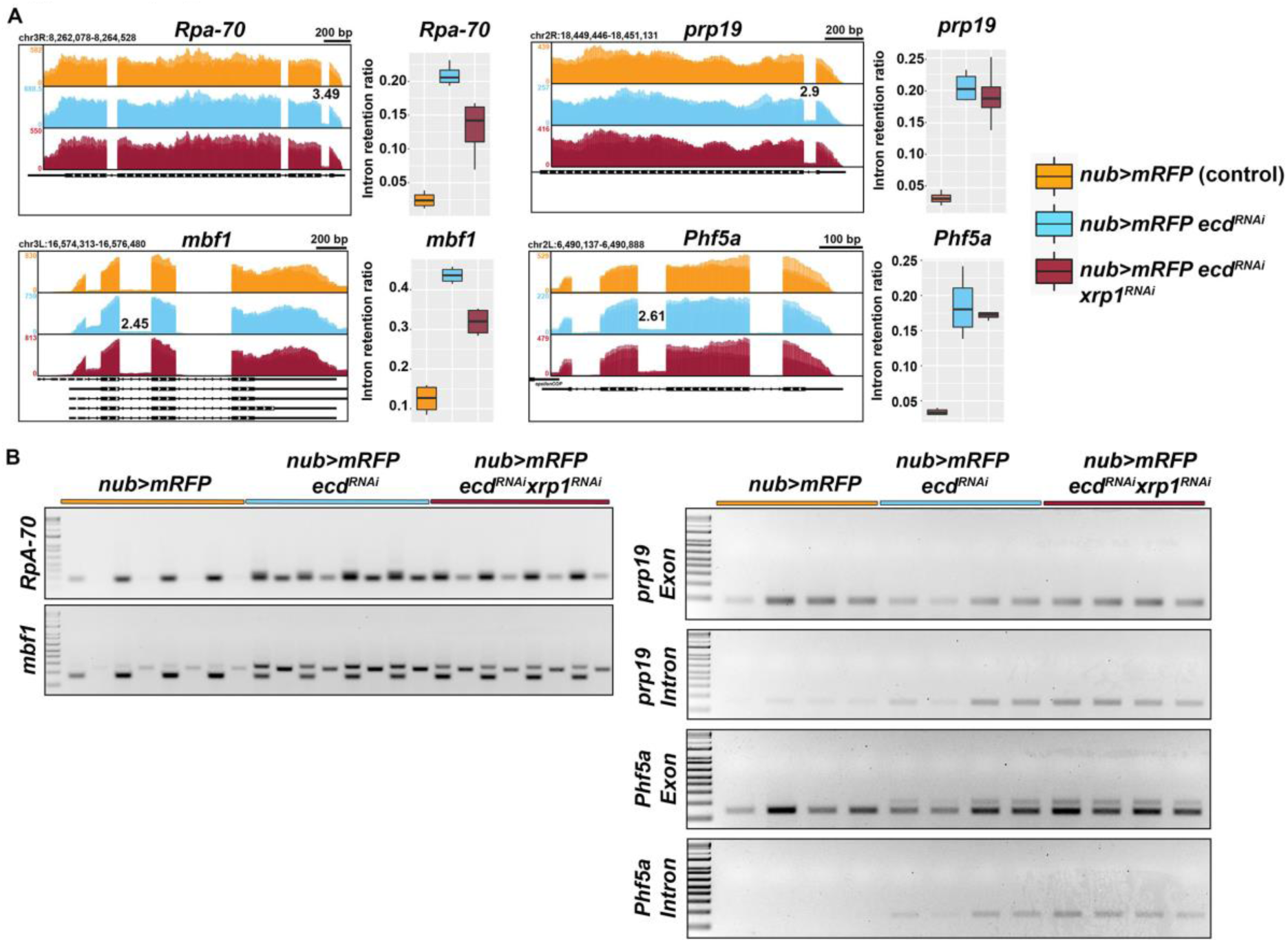
Expression levels but not missplicing are normalized by reducing Xrp1 in Ecd deficient cells. **(A)** Superimposed RNA sequencing coverage of sample replicates across candidate genes bodies with indicated log_2_ Fold Changes above the retained intron (left panels) and intron retention ratios as determined by NxtIRFcore/IRFinder analysis (see Materials and Methods) for the same candidates (right panels). **(B)** Representative gel image of semi-quantitative PCR results using intron and exon-exon junction specific primer sets on RNA samples independent of those used for RNA-seq (n = 4 per genotype) shows the intron retention across replicates in all experimental groups. Exon- and intron-specific amplicons are resolved on adjacent lanes, beginning with an exon for each of the samples. Related to Figure 6. See also Supplementary Dataset S3 and S4.

**Supplementary Table S1.**
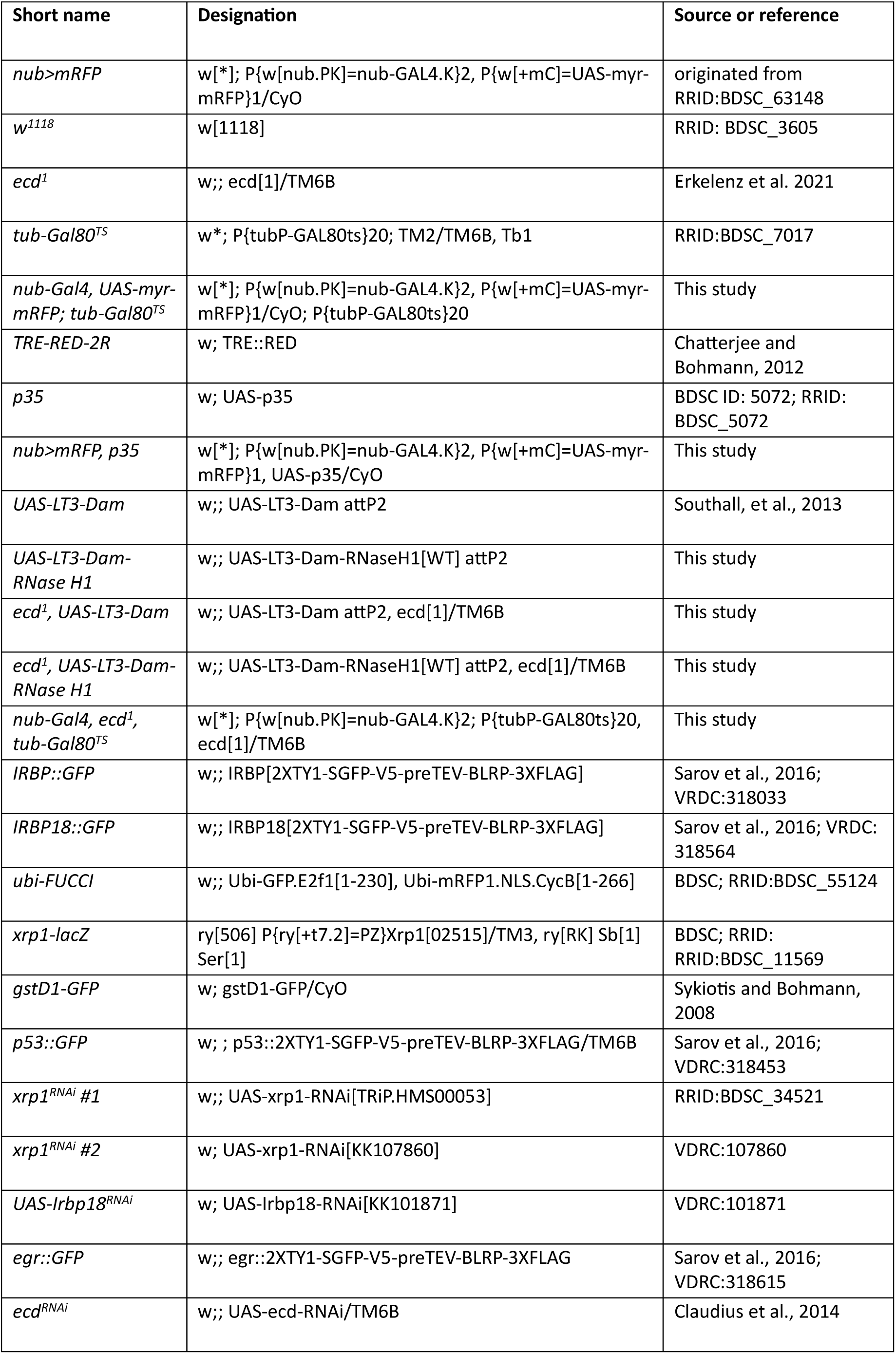
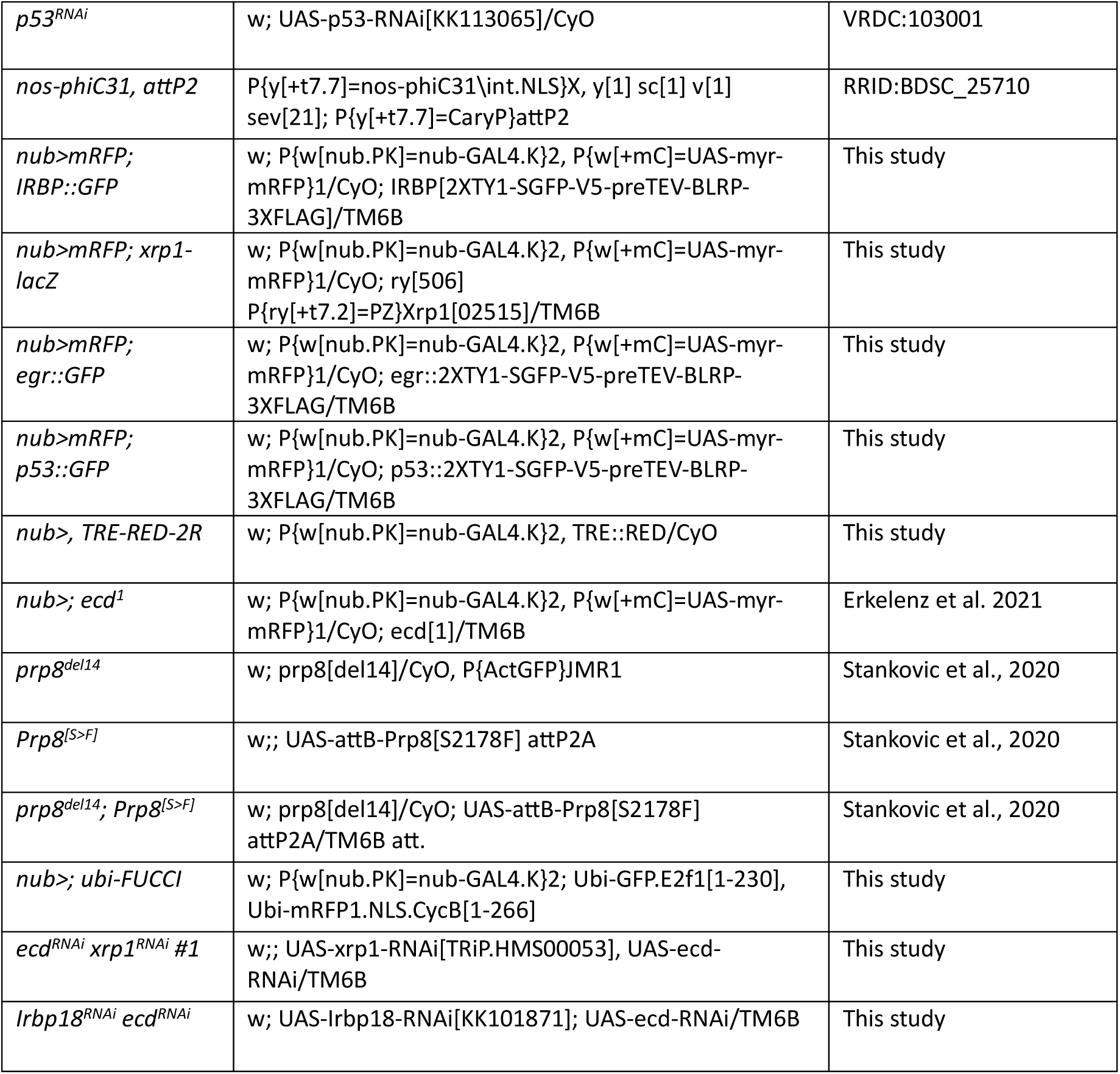
*Drosophila* lines.

**Supplementary Table S2.**
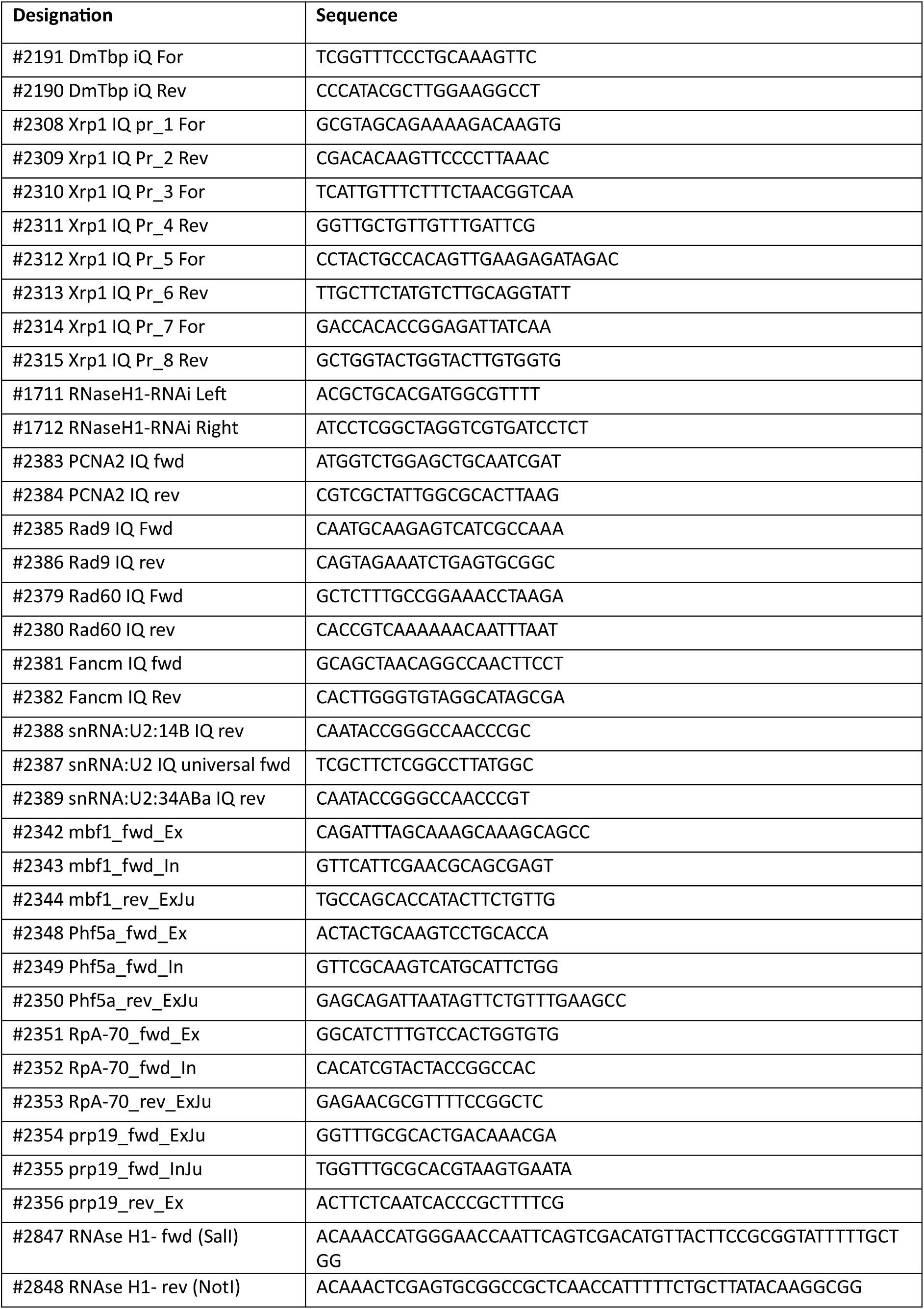
Primers and oligos.

Supplementary Dataset S1. RNase H1 TaDa-seq results.

Supplementary Dataset S2. RNA-seq results (differential gene expression and GO term enrichment analysis) for control (*nub>mRFP*) and Ecd deficient (*nub>mRFP ecd^RNAi^*) wing imaginal discs.

Supplementary Dataset S3. RNA-seq results (differential gene expression and GO term enrichment analysis) for control (*nub>mRFP*), Ecd deficient (*nub>mRFP ecd^RNAi^*), and Ecd, Xrp1 deficient (*nub>mRFP ecd^RNAi^ xrp^RNAi^*) wing imaginal discs.

Supplementary Dataset S4. RNA-seq results (differential splicing analysis, GO term enrichment analysis, and protein-protein interaction network enrichment analysis) for control (*nub>mRFP*), Ecd deficient (*nub>mRFP ecd^RNAi^*), and Ecd, Xrp1 deficient (*nub>mRFP ecd^RNAi^ xrp^RNAi^*) wing imaginal discs.

